# An improved CRISPR-base editor tool to target virulence factors in the ruminant pathogen *Mycoplasma bovis*

**DOI:** 10.64898/2026.05.29.712936

**Authors:** Patrick Hogan, Marc Duclusaud, Thomas Ipoutcha, Carole Lartigue, Geraldine Gourgues, Alain Blanchard, Eric Baranowski, Laure Béven, Yonathan Arfi, Pascal Sirand-Pugnet, Fabien Rideau

**Author notes:** Corresponding author: Pascal Sirand-Pugnet.

## Abstract

*Mycoplasma bovis* is a minimal bacterium infecting cattle, which causes a wide variety of symptoms and is impacting dairy and beef producers worldwide. Part of the difficulty in research surrounding *M. bovis*, and other mycoplasmas, is the lack of efficient genome editing tools. As a proof of concept, we previously presented a transposon-based CRISPR-Base Editor system to introduce targeted mutations in *M. bovis*. In this work, the existing tool has been greatly improved: multi-loci targeting through addition of a second guide RNA; increased number of targetable loci by using an engineered Cas9 with AT-rich PAM specificity, and elimination of the CRISPR-Base Editor from the generated mutants through either transposon excision or use of a curable plasmid. We also propose a dedicated bioinformatic tool to identify target sequences in genes of a given genome. This software was applied to demonstrate the potential of our improved tools in *M. bovis* and other mycoplasmas of veterinary and human interest that currently lack genome editing methods.

## INTRODUCTION

The ability to genetically modify bacteria and generate targeted mutants is fundamental to the study of microbial physiology and gene function. Although well-established in model organisms, these approaches remain largely inaccessible for many species. This is particularly true for mycoplasmas, a group of host-associated bacteria (class *Mollicutes*) responsible for chronic infections in humans, wildlife, and economically important livestock^1^. Genetic tools available for these organisms have long been limited to transposons and replicative plasmids, mainly employed for transposon mutant library production, gene complementation or heterologous expression^2^. The lack of efficient homologous recombination systems in most mycoplasma species has severely hindered molecular studies in these bacteria.

In the mid-2000s, the reduced genome size of mycoplasmas, typically under 1 Mbp, led to their use as minimal cell models for pioneering experiments in Synthetic Biology, including genome cloning and assembly using *Saccharomyces cerevisiae* as a cloning platform, and whole genome transplantation to generate a minimal bacterium^3–5^. However, this revolutionary approach which enables sophisticated genome engineering in mycoplasmas remains applicable to only a few species within or related to the so-called *Mycoides* cluster^6–9^. For many other mycoplasmas relevant to both human and animal health, efficient genetic tools are still lacking, even though the use of exogenous recombinases and CRISPR-based systems have recently opened new perspectives ^10–13^.

Among these is *Mycoplasma bovis*, a major pathogen in cattle causing pneumonia in beef herds and mastitis in dairy cattle, which poses a serious threat to bovine-producing countries worldwide^14^. Moreover, *M. bovis* is also emerging as a significant pathogen in bison herds, a fast-growing livestock industry in North America^15^.

Due to the lack of effective curative treatments, infected herds are often quarantined and culled, and existing vaccines tend to show limited or inconsistent efficacy in the field^16^. The most striking example of the impact of *M. bovis* in recent years is the 2017 outbreak in New Zealand, previously considered free of the disease^17^. Rapid spread of the pathogen across multiple cattle-producing regions led to a nationwide eradication campaign involving massive diagnostic testing, pre-emptive culling, and financial compensation of farmers. This initiative—coordinated by the New Zealand government and dairy/beef industry partners—aims to eliminate *M. bovis* by 2028^18^. While it has significantly reduced pathogen prevalence, the program has come at a high cost: over 180,000 cattle culled, considerable psychological stress on farmers, and a projected budget of NZD 886 million (∼400-500 million USD) over 10 years^19^.

Understanding the virulence factors and genetic determinants of *M. bovis* pathogenesis is therefore essential to developing more effective and less disruptive control strategies. In this context, a CRISPR-Base Editor (BE) system adapted in our group was recently demonstrated to be functional and efficient in several veterinary-relevant mycoplasma species, including *M. bovis*^10^.

Originally developed by Komor *et al.*^20^, the CRISPR-BE comprises a catalytically inactive *Streptococcus pyogenes* Cas9 (SpdCas9) fused to the cytosine deaminase pmcDA1 and an Uracil DNA Glycosylase Inhibitor (UGI). Upon gRNA binding, the complex recognizes the NGG Protospacer Adjacent Motif (PAM) site and induces local DNA duplex opening at the target locus. While the endonuclease domains are inactivated (preventing double-strand breaks), exposed cytosines are converted to uracil by pmcDA1, leading to C→T transitions (G→A on the complementary strand) during replication. UGI blocks the base excision repair pathway, ensuring mutation fixation.

This original CRISPR-BE tool was adapted to mycoplasmas by recoding its coding sequence to accommodate the AT-rich genomes of mycoplasmas (24–31% GC content) and changing the promoters controlling the expression of the gRNA and the SpdCas9-pmcDA1-UGI. The CRISPR-BE tool components are carried by a pMT85/2res plasmid derived from the Tn*4001* transposon^21^. A proof-of-concept experiment in *M. bovis* was performed and demonstrated the ability to introduce a premature stop codon in *mnuA*, a gene encoding a membrane nuclease involved in degrading neutrophil extracellular traps (NETs) during infection^10^. While efficient (tens of mutant clones obtained for each transformation and induction cycles), this system also presents several limitations.

First, the original tool includes one single gRNA which might be limiting for engineering plans involving several modifications. Second is the PAM limitation, as the SpCas9 enzyme requires an “NGG” PAM, which is relatively scarce in the AT-rich genomes of mycoplasmas. This limits the number of editable cytosines and, consequently, the possibility to generate stop codons and inactivate genes. The third is the Transposon-Based Delivery, as the BE system is carried on a transposon whose random integration can lead to unintended gene disruption or dysregulation, complicating interpretation of phenotypes. The final limitation is the permanent integration of the CRISPR-BE genetic tool, including an antibiotic resistance marker (gentamicin), which precludes its reuse in iterative mutagenesis. In addition, the constitutive presence of the CRISPR-BE module, even in the absence of inducer, increases the risk of off-target mutagenesis through observed P*xyl/tetO(2)* promoter leakage^10^.

To overcome these limitations, we constructed and tested a new suite of CRISPR-Base Editing tools tailored for iterative, high-precision, and scarless genome editing in mycoplasmas. These improvements include the ability to perform simultaneous multi-targeting using a single plasmid to express a dual gRNA allowing multiple knockouts of distinct genes; as well as an expanded targeting range through the use of an engineered SpdCas9 variant that recognizes the NGA PAM, thereby increasing target site availability in AT-rich genomes. We also provide a strategy to remove the editing cassettes, with a post-editing excision of the BE module and antibiotic marker via site-specific recombination. We then transferred these improved systems to a replicative and curable plasmid backbone for transient editing, enabling clean genome modifications without residual foreign DNA. Finally, we propose a dedicated bioinformatic tool that can be used to identify suitable NGG/NGA targets in any target genome.

These improved tools are used to perform the targeted inactivation of several candidate virulence genes in *M. bovis*, paving the way for accelerated functional studies and the development of next-generation control strategies.

## RESULTS

The CRISPR-BE tool we have previously developed (pMT85_SpdCas9_pmcDA1^10^, here named as pFRIT4.0) is based on a transposon, which carries the complete edition tool and a selection marker (Fig. S1). After growth on selective media, the transformants are isolated and passaged in liquid media, followed by induction of the expression of the SpdCas9-pmcDA1-UGI by addition of anhydrotetracyline (aTc). Meanwhile, the single gRNA including the 20 nt target sequence and the tracrRNA structural part is expressed constitutively under the control of the spiralin promoter and fibril terminator. Following one or more rounds of induction, the edition of the target locus is screened by Sanger sequencing ^10^.

### Producing multigene knockouts in *Mycoplasma bovis*

Our first goal was to expand the ability of our system to target multiple loci during one single mutagenesis process. To do so, we added a second gRNA expression cassette to our integrative vector (Fig. 1A). This second cassette is identical to the first one, with the exception of the target sequence spacer.

**Figure 1.**
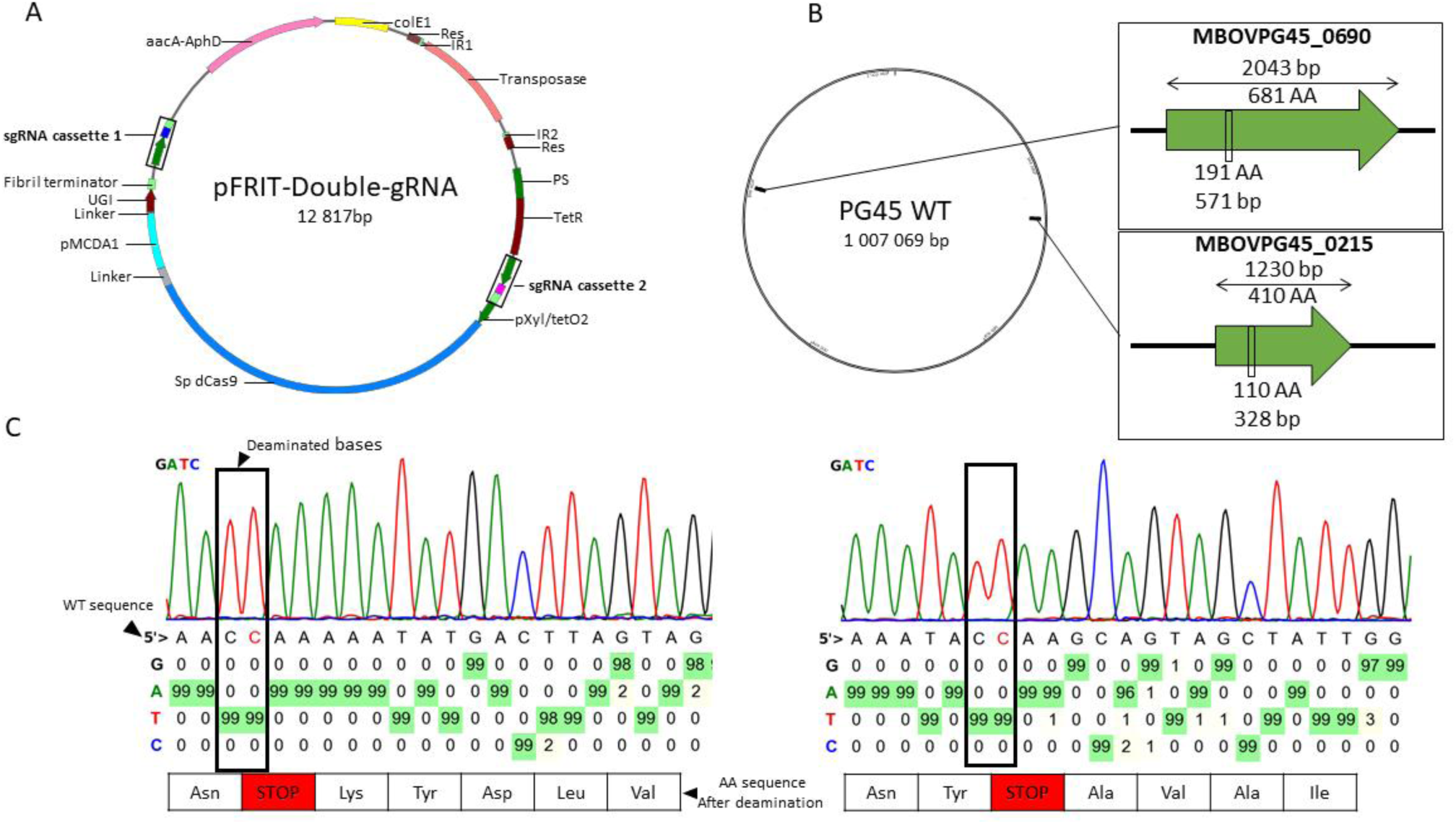
Simultaneous targeting of the *mnuA* and *5’ nucleotidase* genes. **(A)** pFRIT4.0-DgRNA plasmid map. The two sgRNA cassettes are indicated **(B)** Genome positions of MBOVPG45_0215 (*mnuA*) and MBOVPG45_0690 (5’-*nt*) targeted genes and respective location of the generated stop codons within genes. **(C)** Sanger sequencings of the targeted sites within MBOVPG45_0215 (left) and MBOVPG45_0690 (right) of an isolated mutant. The percentage of bases found in the isolated clones of transformants were determined from Sanger sequencing chromatograms using the Base editor processing tool from Han lab^23^.

To benchmark the ability of this dual-target CRISPR-BE tool in *M. bovis*, we attempted the simultaneous editing of two distinct genes located at opposite points of the genome (Fig. 1B). The first locus MBOVPG45_0215 encodes the nuclease MnuA and was previously successfully targeted^10^. The second locus, MBOVPG45_0690, encodes a 5’ nucleotidase (5’-NT) and was shown to be implicated in fitness and virulence of *M. bovis* in mastitis cases^22^. To construct the double target plasmid, we took the previously validated single gRNA targeting MBOVPG45_0215 plasmid and cloned a second empty gRNA cassette (without an inserted target sequence) within the plasmid. The target sequence for MBOVPG45_0690 was then cloned into the new empty gRNA cassette through a digestion-ligation method. The two gRNAs were designed to introduce mutations and transform codons encoding Q191 (MBOVPG45_0215) and Q110 (MBOVPG45_0690) into stop codons (Fig. 1B). The constructed plasmid (Fig. 1A) was then used to transform *M. bovis* PG45 cells. After transformation, cultures were induced overnight and plated on selective media. Eight clones were isolated for screening by PCR amplification and sequencing of the target sites. Within the eight screened transformants, three were fully deaminated on the first screened locus (*mnuA*, MBOVPG45_0215). Among those, one mutant was also fully deaminated on the second screened region (5’-NT encoding gene MBOVPG45_0690) (Fig. 1C). The two other *mnuA* mutants revealed partial deamination of the second gene. This experiment demonstrated the efficiency of the dual-gRNA CRISPR-BE to target two different genes through a single round of induction and the screening of a limited number of transformants.

Following this successful targeting of two genes, we then tried to expand this approach to simultaneously disrupt a family of genes using two other target sequences. Specifically, we targeted the transposable IS*Mbov1* insertion sequences, which are a part of a diverse set of mobile genetic elements present in *M. bovis*. To date, seven families have been characterised in this species (IS*Mbov1* to IS*Mbov7*)^24,25^. Within the *M. bovis* PG45 type strain, 11 copies of IS*Mbov1* were identified after whole genome sequencing (Fig. 2A). Although the nucleotide sequences of IS*Mbov1* copies are highly conserved (>97%), a single gRNA targeting all copies could not be designed. Two regions were then targeted within IS*Mbov1* sequences to introduce stop codons and generate premature truncation of the encoded transposase at two positions, Q108 (Target 1) and Q212 (Target 2) (Fig. 2B).

**Figure 2.**
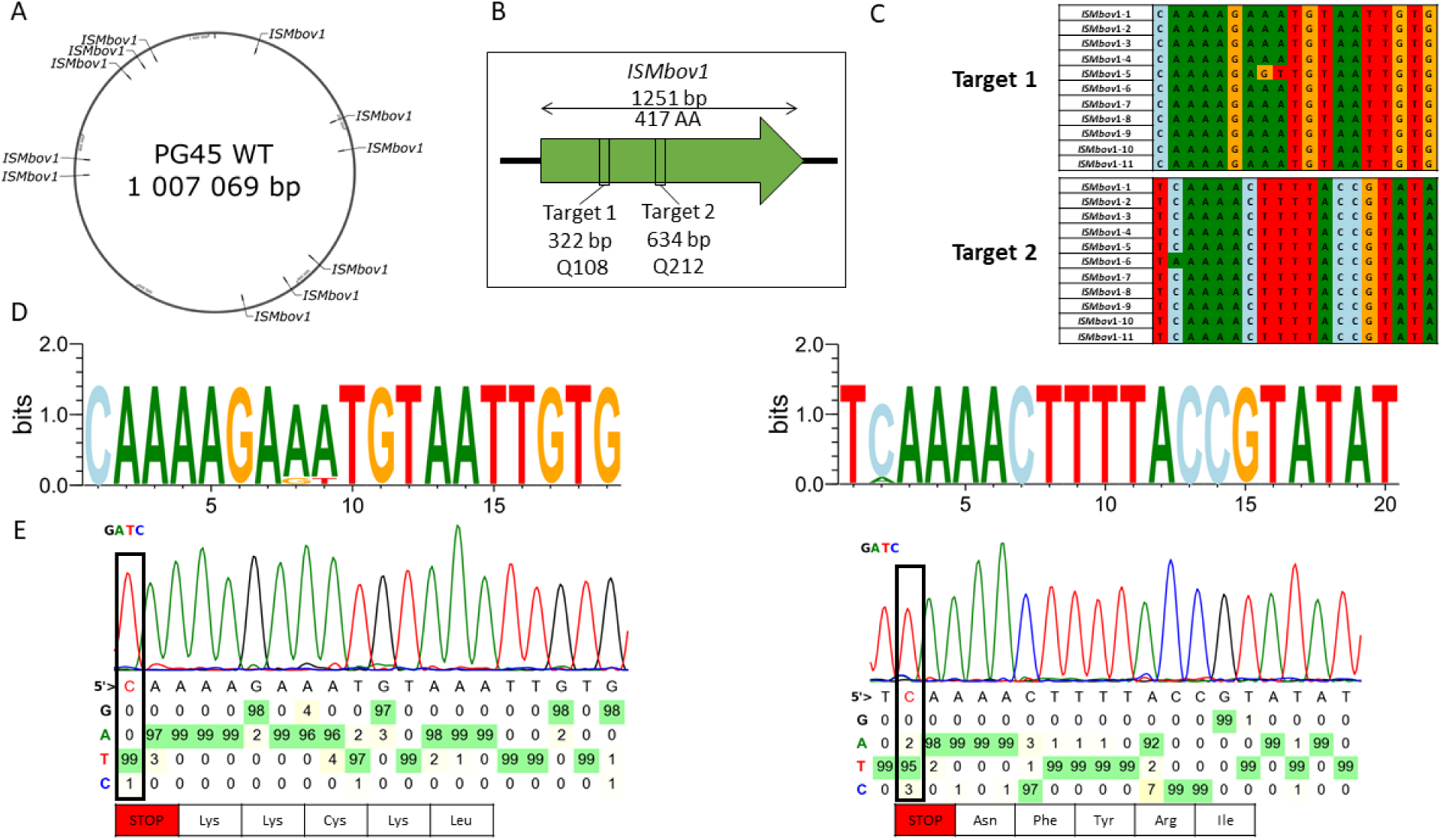
Simultaneous knockout of all IS*Mbov1* copies in *M. bovis* PG45. **(A)** Localisation of the 11 copies of IS*Mbov1* within the genome of *M. bovis* PG45 strain. **(B)** Targeted regions of IS*Mbov1* copies through double gRNA expression and the respective location of the generated stop codons **(C)** IS*Mbov1* isoform sequence conservation of targeted regions (representation through muscle alignment). **(D)** gRNAs target sequences conservation within IS*Mbov1* done through Web-logo^26^. **(E)** Sanger sequencing of the targeted sites, Target 1 (left) and Target 2 (right) within an isolated mutant after three rounds of induction. Chromatograms were analysed using the Base editor processing tool from Han lab^23^.

Each target site is present within 10 out of 11 IS*Mbov1* copies allowing for broad but not complete coverage (Fig. 2C and 2D). The dedicated DgRNA-pFRIT plasmid was built through an intermediate construction of two single gRNA pFRIT plasmids that were then used as templates for a DgRNA assembly (detailed in materials and methods section). After transformation and plating on selective medium, three colonies were picked and cells were grown selective liquid medium before overnight induction. Following induction, the three populations of cells were screened through PCR amplification and Sanger sequencing on both targeted regions. All three presented partial deamination profiles suggesting a single round of induction was insufficient, potentially due to the number and distribution of targets, to obtain fully deaminated mutants. However, after two additional induction cycles, populations of cells showing improved deamination profiles on both target sites were obtained for the three clones, with one clone showing nearly complete deamination (Fig. 2E and S3).

### Expanding the repertoire of targetable genes using a VQR variant of SpdCas9

One of the critical steps in the deamination by the CRISPR-BE tool is the recognition of the PAM sequence. However, in the case of bacteria of the class *Mollicutes*, owing to the low G+C content of their genomes, the NGG PAM sequences recognized by SpCas9 are relatively scarce. Analysis of the 766 coding sequences of *M. bovis* PG45 reveals a total of 19,066 NGG nucleotides triplets. For a successful truncation of the CDS, these PAM are also required to be downstream of a cytosine, which needs to be within a 15-21 bp editing window and for which the conversion into uracil will generate a stop codon. Therefore, out of the 19,066 NGG, only 1,475 are of interest here. To expand the number of potential target sites, alternative SpCas9 variants with different PAM recognition specificities could be employed. One such variant is the VQR-SpCas9, formed from the substitution of three specific amino acids D1135V/R1335Q/T1337R, allowing the recognition of a NGA PAM sequence^27^. A total of 54,163 NGA sequences are present in the coding sequences of *M. bovis* PG45, including 3,975 of interest, a roughly 2.7-fold increase compared to the original NGG PAM. The VQR variant CRISPR-BE tool for mollicutes (pFRIT-VQR plasmid) was derived from the original single gRNA tool (pFRIT4.0 plasmid), through the substitution of four nucleotides and leading to the three necessary amino acid changes. To evaluate the efficiency of this new system, we targeted independently two genes that could not be disrupted by the original version of the tool. These genes, MBOVPG45_0373 and MBOVPG45_0376, encode two Mycoplasma Immunoglobulin Proteases (MIP) implicated in the capture and cleavage of host immunoglobulins in the MIB-MIP system^28^. The targeted sites were selected to introduce premature stop codons corresponding to Q228 (MIP-1, MBOVPG45_0373) and Q66 (MIP-2, MBOVPG45_0376) (Fig. 3A), respectively. Following the molecular cloning of the target sequences into the gRNA cassettes, the two generated single gRNA plasmids were transformed into mycoplasma cells to inactivate MBOVPG45_0373 or MBOVPG45_0376. For each target, five transformants selected on gentamycin plates were picked and cultures for three passages before an ON induction with aTC. Three clones for each target were analyzed by PCR amplification and sequencing of the targeted genome sites. Clones showing the best deamination profiles were selected and submitted to another round of induction before filtration and plating on gentamycin plates. For each target, one subclone was selected for analysis by PCR and sequencing, showing the expected mutation (Fig. 3B).

**Figure 3.**
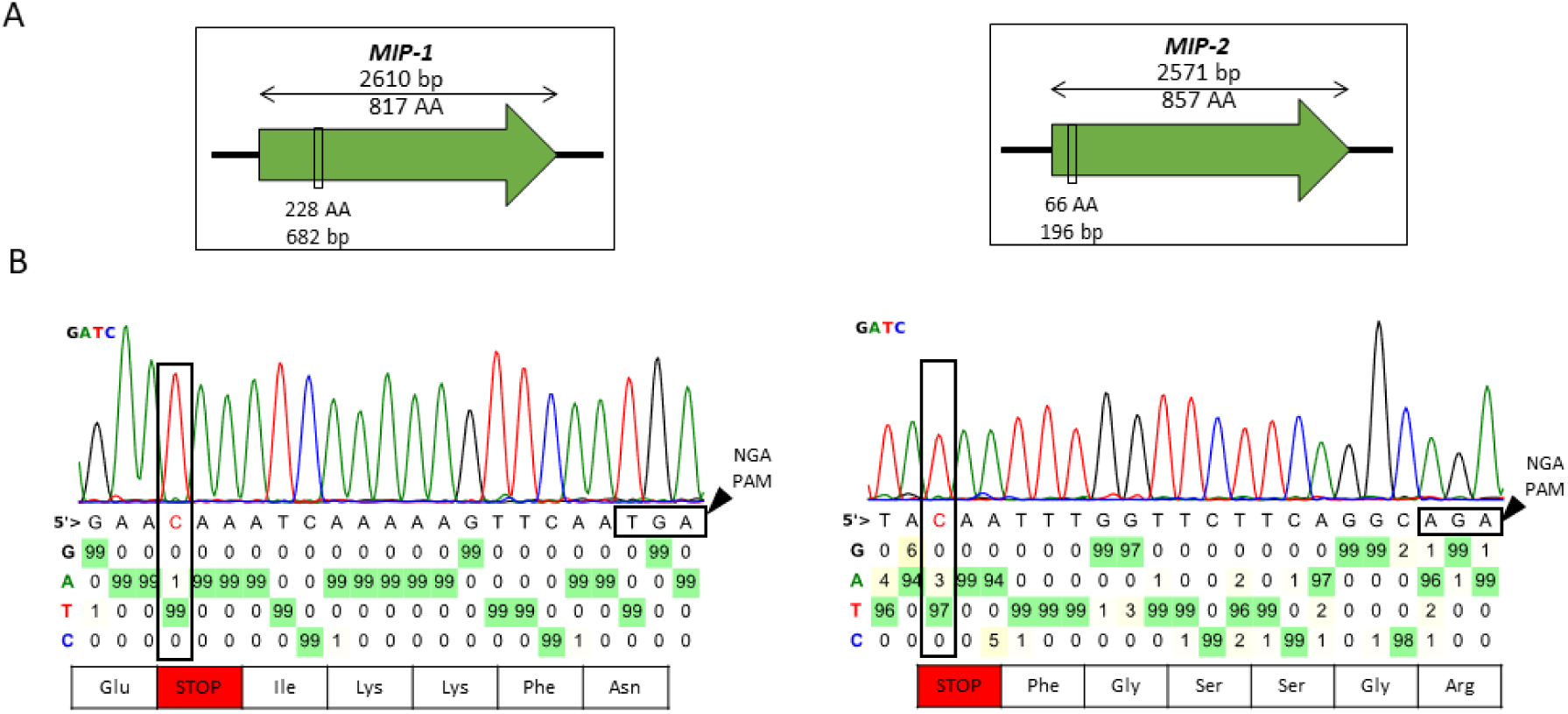
Targeting MIP genes with the VQR variant of SpdCas9. **(A)** Targeted regions of MIP genes conducted with CRISPR-VQR variants. **(B)** Representation of Sanger sequencing of the targeted sites within MBOVPG45_0373 (left) or MBOVPG45_0376 (Right) within two isolated mutants. The black square indicates the used PAM sequence. Sanger sequencing chromatogram analysis was done using Base editor processing tool from Han lab^23^.

### Eliminating the CRISPR-BE tool after successful knockouts

The possibility to remove the integrated CRISPR-BE tool from the genome after deamination is desirable for several reasons. First, this would limit the accumulation of unwanted off-target mutations potentially generated by the active editing system. Second, the elimination of the system and the associated antibiotic resistance marker would open the way to iterative mutagenesis. Third, engineered bacterial strains cleaned of genetic tools and markers are often required for applied uses as vaccines. Therefore, we elected to develop a strategy to eliminate the CRISPR-BE tool after mutagenesis.

The original CRISPR-BE tool was constructed using the pMT85/2res backbone as a vector^21^. This plasmid harbours a transposon and two *res* resolution sequences that enable excision of the integrated transposable element through the use of a secondary replicative plasmid containing the γδ resolvase gene (*tnpR*). The recombination between the *res* sequences under the action of the resolvase only leaves a minimal scar of 218 bp in the genome, corresponding to a single *res* site flanked by the two inverted repeats (Fig. S4). To carry out the resolution, we constructed the pRES plasmid (Fig. S5), based on the original pCJ15 replicative plasmid^21^, by replacing the replicative origin of *Mycoplasma mycoides* subsp*. mycoides* by that of *Mycoplasma agalactiae*, which is proven to be functional in *M. bovis* ^29^.

To test the functionality of pRES, the double knockout mutant MBOVPG45_0215-MBOVPG45_0690 obtained in this study (Fig. 1) was subsequently transformed with the pRES plasmid. Ten transformants were picked on tetracycline-supplemented plates and further cultured for two passages in presence of tetracycline (Fig. S6 and detailed in the Materials and Methods section). Excision of the transposon harbouring the CRISPR-BE and the gentamicin resistance marker was monitored through PCR amplification of the resolution site and loss of the ability to grow in gentamicin selective media. One additional step of subcloning was necessary to isolate resolved clones.

Two selected resolved clones were then cultivated under non-selective conditions to cure *M. bovis* mutants from the pRES plasmid. After 10 passages and one subcloning step, the pRES elimination was confirmed by growth inhibition on tetracycline selective media. Finally, the presence of the intended mutation was reconfirmed through PCR amplification, Sanger sequencing and a final verification of resolution was performed through whole genome sequencing (Fig. S7).

Although the use of the pRES plasmid to drive the excision of the CRISPR-BE and the selection marker was a success, the use of an integrative plasmid still places limits on further applications of generated mutants. Indeed, the insertion locus of the transposon harbouring the CRISPR-BE and the selection marker is random and elimination of the integrated cassette by resolution leaves a scar that may have unwanted impacts on the engineered strains. Any generated truncation or gene modification is thereby conserved regardless of resolution. For instance, whole genome sequencing of the resolved MBOVPG45_0215-MBOVPG45_0690 double mutant revealed that the transposon was inserted at 813 pb within the MBOVPG45_0756 coding sequence, coding for a putative hydrolase, truncating the gene prematurely (Fig. S7). Even in case of intergenic insertion of the transposon, unexpected impact of the remaining scar may be difficult to anticipate. In addition, the complete process involves two distinct transformations and up to twenty passages under varying selective and non-selective conditions. While effective, this workflow is time-consuming and subjects the mutant populations to repeated stress events. Furthermore, excessive passages increase the likelihood of divergence from the original transformed mutant due to single nucleotide polymorphisms (SNPs), insertions/deletions (indels), or mobilization of genetic elements.

For these reasons, we elected to also develop a set of CRISPR-BE tools for *M. bovis* based on a replicative backbone, such as already described for *Mycoplasma mycoides* subsp*. mycoides*^10^. Once deamination is successful, these plasmids can be eliminated rapidly through passages in a non-selective medium (Fig. S8). We built two replicative plasmids, called pFrHog and pFrHog-VQR (Fig. S9), based on the backbone p20-1miniO/T^30,31^ carrying the *oriC* from *M. agalactiae*. Into this backbone, we assembled the gentamicin resistance marker with either the CRISPR-BE system or with the CRISPR-BE VQR system from our pMT85/2res-based plasmids.

As a proof of concept, we targeted MBOVPG45_0215 (*mnuA*) with the same gRNA as previously described. After transformation, induction and plating on gentamicin selective media, six out of six screened transformants displayed the desired mutation. Two transformants were then chosen at random, cultured in non-selective medium for two additional passages and subcloned on non-selective plates for curing of the replicative plasmid. Three plated subclones of each culture were then picked and grown for three additional passages in non-selective media. Curing confirmation was conducted through PCR verification and failure to grow in selective media.

Overall, we have developed a set of five new plasmids, starting from our original single target Base Editor pFRIT4.0 (Fig. 4). First, the pFRIT4.0-DgRNA enables multi-target editing. Meanwhile, the pFRIT-VQR offers an extend target range through recognition of an NGA PAM. Its derivative pFRIT-VQR-DgRNA introduces multi-targeting to the VQR variant. Finally, pFrHog and its derivative pFrHog-VQR enable single target editing in an easily removable vector. This set of tools delivers a drastic improvement to the capabilities of CRISPR-BE tools in *M. bovis*, enabling the rapid generation of new *M. bovis* mutants and paving the way towards a better understanding of this pathogen’s biology.

**Figure 4.**
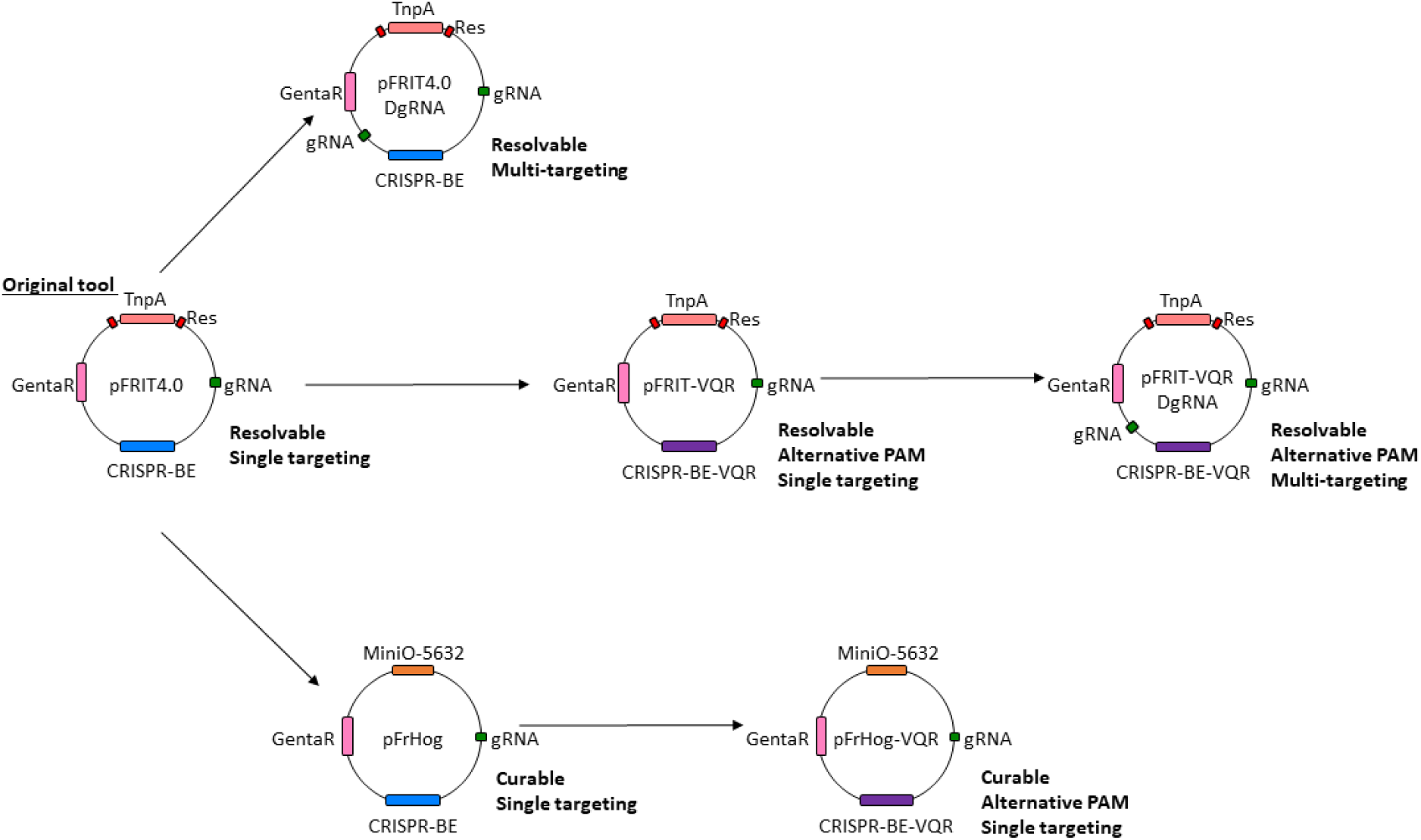
Extended repertoire of CRISPR-BE tools validated for use in *M. bovis.* Schematic representation of the various CRISPR-BE plasmids developed and used in this work. All plasmids were based on the original pFRIT4.0 plasmid^10^.

### Genome-scale mapping of target sites for CRISPR-BE and CRISPR-BE VQR tools

To get a broader view of the genes targetable for knockout by our Base Editor tools, and facilitate the design of the corresponding gRNAs, we have created a Python-based algorithm, called Deaminase Target Identifier (DTI) (Fig. S10). Based on a user-defined genome, DTI identifies and presents all the potential target sequences that could be used to generate stop codons in coding sequences. The main flow of DTI is as follows. First, the user inputs two fasta files containing the target genome and the target CDSs. Then, a set of variables are selected to modulate the search preferences such as the targetable window size and preferred PAM sequence in proximity to the targeted codon. Based on the start codon of each submitted coding sequence, the algorithm searches along the ORFs for sequences that could be targeted to generate stop codons. They can be generated by targeting CAA/CAG glutamine codons to convert them into TAA/TAG stop codons, or by targeting a TGA tryptophan codon through deamination of the complementary C on the reverse strand to generate a TAA stop codon. Once one of these codons of interest has been identified, the presence of an upstream PAM sequence within the chosen editing window is evaluated. The algorithm then outputs a list of all the possible target codons and the associated target sequence.

Using this tool, we have evaluated and compared the potential of the CRISPR-BE and CRISPR-BE VQR tools for gene knockouts in *M. bovis* PG45. In order to inactivate genes, the position of the generated stop codon is crucial, with earlier truncation potentially providing functional knockouts whereas later positions may only reduce functionality or have no impact at all. We therefore categorized all the generable stop codons by their relative position in the CDS: either within the first 10% of the CDS length, the first 30% or the first 50%. We also included two alternative categories: truncation at any point beyond 50%, and non-targetable (Fig. 5).

**Figure 5.**
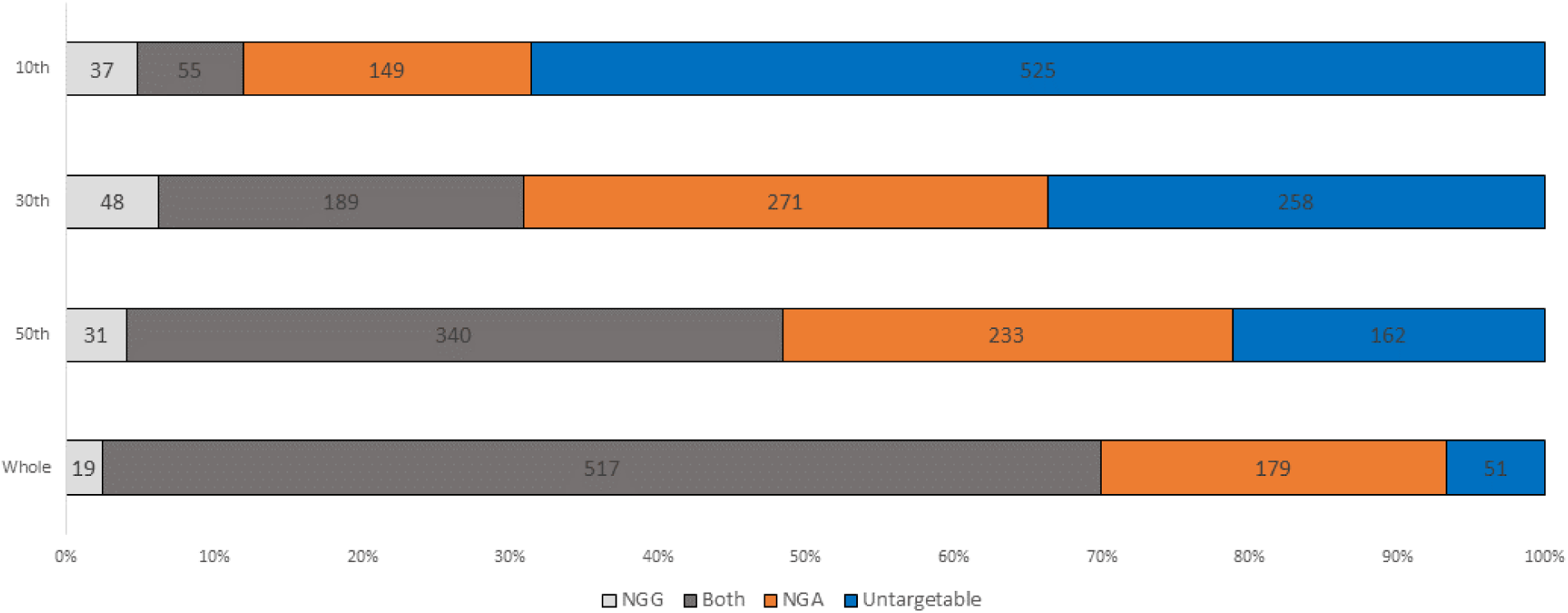
Distribution of gene truncation potential by developed CRISPR-BE tools. Distribution of the 766 *M. bovis* PG45 coding sequences categorised by their targetability by CRISPR-BE tools within the 10^th^, 30^th^, 50^th^ and whole of gene length. Targetable genes are then subcategorised by the PAM found in proximity to the targeted codon, NGG - Light grey; NGA-orange; Both-dark grey. Finally untargetable genes are in blue.

Based on these categories, we calculated that out of the 766 annotated CDS present in the *M. bovis* PG45 genome a total of 715 were targetable by using either of the developed BE tools. Of these, 604 could be targeted within the first half of the coding sequence, 508 within the first third, and 241 within the first tenth (Fig. 5). As expected, the number of targetable genes decreased steadily when the possible position for stop introduction became smaller.

In-line with the increased number of NGA sequences in coding sequences compared to NGG, the number of generatable stop codons in the first 10^th^ and 30^th^ using the CRISPR-BE VQR tool is doubled compared to that for the original CRISPR-BE tool. This observation further highlights the new possibilities offered by the VQR variant for targeted knockouts in genes that were previously inaccessible. However, it is also noteworthy that the VQR variant does not fully replace the original tool, as a limited number of genes are only targetable through the original NGG PAM. As the stringency for an early stop decreases, the overlap between both tools increases, indicating that both tools could be used for targeted gene inactivation. In these cases, the choice of which will be driven by the desired truncation position. Overall, these results demonstrate that both tools are complementary, depending on the gene of interest and the preferred location of deamination.

To further evaluate the potential of our CRISPR-BE tools across multiple species of mycoplasma, we employed the DTI algorithm on a set of veterinary and human relevant mycoplasma genomes, many of which currently lack efficient genome engineering possibilities. As for *M. bovis*, we have tested the ability of either CRISPR-BE or CRISPR-BE VQR to introduce stop codons within all the CDS predicted in the genomes. Our results indicate that for most mycoplasma species, 27-41% of genes can be targeted within the first 10^th^ of CDS length (Fig. 6). The major outlier is *M. mycoides* subsp*. mycoides* T1/44, in which 63% of genes can be truncated at this level. From then on, roughly 80-90% of all genes can be theoretically truncated at half of gene length. These findings suggest that the CRISPR-BE and CRISPR-BE VQR tools developed here could be effective across many mycoplasma species, enabling the generation of numerous targeted knockout mutants.

**Figure 6.**
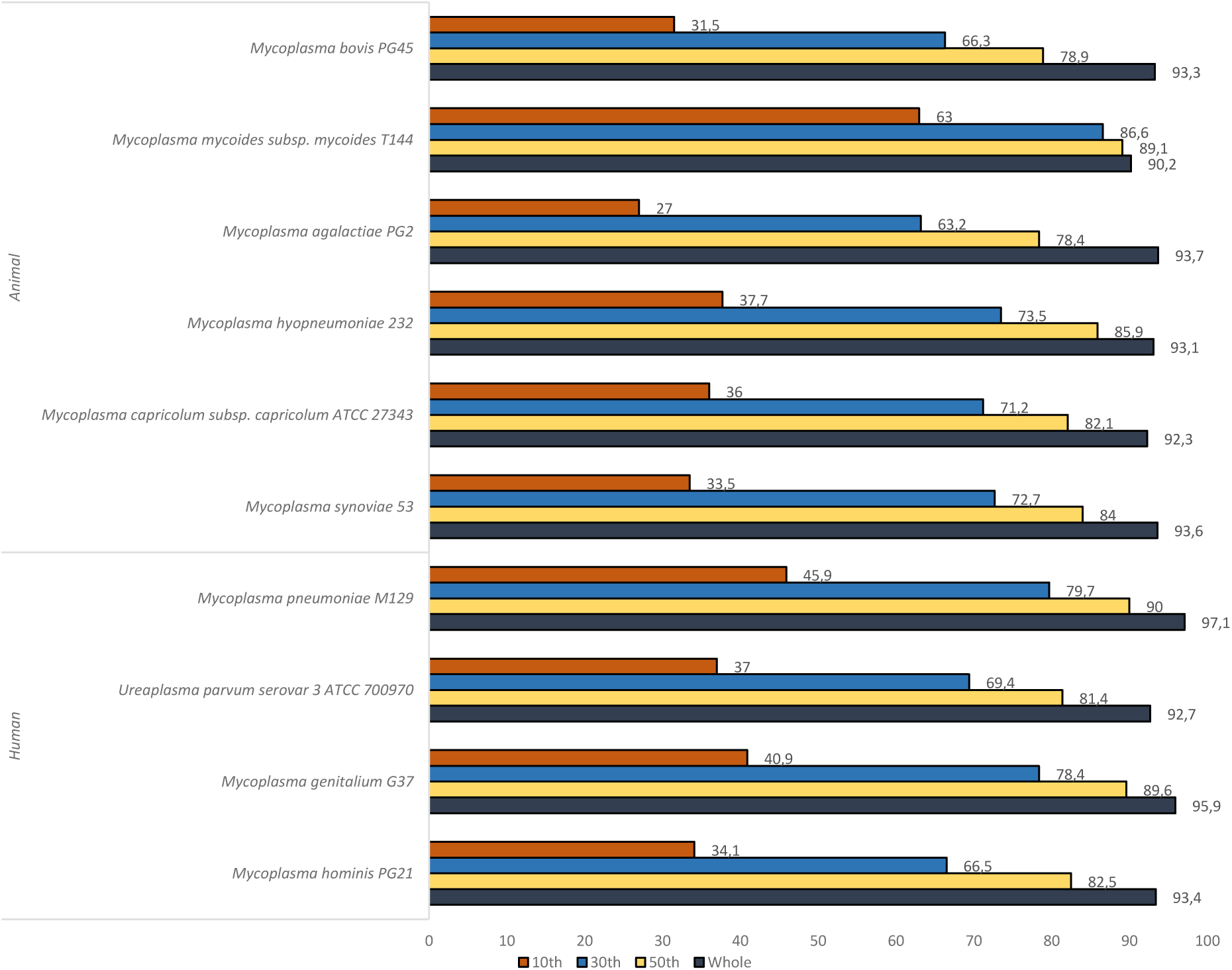
Predicted potential of CRISPR-BE to target genes of interest in human and animal relevant mycoplasma species. Representation of the percentage of targetable CDS by the CRISPR-BE or CRISPR-BE VQR tools compared to the total number of CDS in multiple mycoplasma species. Representation of the percentage of genes targetable within the first 10^th^, 30^th^, 50^th^ and whole coding sequence.

## DISCUSSION

The development of advanced genomic tools is essential to accelerate pathogen research and inform effective disease control strategies. In the face of increasing pathogen emergence and re-emergence, global health risks are intensifying, driven by climate change and globalization, which enhance pathogen dissemination. Rapid identification of the key determinants of host–pathogen interactions has therefore become a critical challenge across human, animal, and plant health.

In this perspective, our team is developing innovative genetic tools for mollicutes, with a particular focus on veterinary mycoplasmas such as *M. bovis*. Recently, we described a first generation of CRISPR-Base Editor tools that proved effective in three major pathogens affecting poultry (*Mycoplasma gallisepticum*) and cattle (*M. mycoides* subsp. *mycoides* and *M. bovis*)^10^. Practical use of these tools has also highlighted their limitations, as some genes could not be inactivated due to the dependency of the original SpdCas9 on an NGG PAM. Moreover, mutant generation was labour-intensive. Thus, improved tools were required to expand targetability, enable iterative cycles of mutagenesis, and reduce turnaround times.

In this work, we expanded the toolbox of mutagenesis methods available in *M. bovis* with novel CRISPR-Base Editor variants designed to meet these needs. First, we demonstrated that it was possible to efficiently inactivate two *M. bovis* genes simultaneously by introducing a second gRNA in an all-in-one vector. The feasibility of multiplex gene inactivation by gRNA co-expression has already been demonstrated in other bacteria^32–36^. The number of gRNAs and the design of their expression cassettes are key determinants of such multi-targeting strategies. One strategy, which we employed, involves combining several monocistronic gRNA expression cassettes. For instance, the MACBETH method, enabled single-, double-, and triple-locus editing in *Corynebacterium glutamicum* with base editing efficiencies of 100%, 87.2%, and 23.3%, respectively^36^. Similarly, in *Shewanella oneidensis*, this strategy enabled simultaneous inactivation of 3, 5, and 8 genes, with reported efficiencies of 83.3%, 100%, and 12.5%, respectively^34^. While efficiency globally decreases with an increasing number of targets, it remains sufficiently high to allow the screening of multi-gene mutants with limited efforts. An alternative approach relies on polycistronic expression cassettes, where multiple gRNAs are expressed under a single promoter. This design is more complex, as it requires the addition of internal cleavage sequences for transcript maturation and release of functional gRNAs. In *Shewanella oneidensis*, this strategy proved less effective than the monocistronic approach^34^. Meanwhile a polycistronic strategy using Csy4 endoribonuclease recognition sites was successfully developed for *Streptomyces coelicolor* (CRISPR-mcBEST), allowing simultaneous inactivation of 17 out of 28 targeted genes^35^. However, considering cloning complexity and reduced efficiency, the authors recommended a maximum of nine gRNAs per cassette. In our study, we demonstrated that simultaneous inactivation of two genes in *M. bovis* was highly effective, with a single induction being enough for isolating three fully deaminated mutants within the first eight screened mutants for the double *mnuA* and 5’-NT gene targeting. In the future, designing polycistronic cassettes for mycoplasmas could represent a further improvement of the tools we have developed, though this would require adapting an effective gRNA maturation system for these bacteria.

Then, to broaden targetability, we have developed CRISPR-BE variants, in which the Cas9 PAM-interacting domain (PI) was mutagenized to alter PAM specificity. Mutations were selected based on the work of Kleinstiver and colleagues who demonstrated that VQR mutations could change PAM recognition in zebrafish embryos and human cells^27^. In mycoplasmas, this VQR CRISPR-BE variant was also effective, as with an NGA PAM preference, this variant is particularly advantageous given the AT-rich genomic bias of these bacteria. Future developments in genome engineering of mollicutes will benefit from the ever-increasing range of Cas9 variants or other enzymes derived from CRISPR systems. For example, some PAM-less Cas9 variants have been shown efficient in various organisms, nearly unlocking entire genomes for precision editing ^37,38^.

Beyond expanding the targeting possibilities of editing tools, functional studies and applied genome engineering often require iterative mutant construction, enabling detailed analysis of each targeted gene’s role. We therefore investigated the possibility of removing the CRISPR-BE cassette and the associated selection marker harboured by the transposon vector after generating a first mutant. Our initial approach relied on previous work using transposon excision by site-specific recombination between two *res* sequences mediated by the γδ transposon resolvase ^21,39^. This strategy proved functional in *M. bovis*, and several resolved mutants were obtained. However, excision required an additional transformation step, followed by plasmid curing in non-selective medium, which prolonged the process and potentially favoured genetic drift. Furthermore, excision leaves behind a residual scar of 218 bp containing inverted repeats (from the transposon) and a *res* site. Thus, although both the selection marker and CRISPR-BE cassette are removed—enabling further mutagenesis cycles—the resulting genome is not scarless, and this scar may have unpredictable effects. To address this limitation, we developed a replicative plasmid-borne CRISPR-BE system. It proved effective, and plasmid curing was achieved after a total of five passages (including sub-cloning) in non-selective medium, drastically reducing the time to mutant generation and leaving no genetic scar. In rare cases, we have observed the spurious integration of the *oriC* plasmid into the *M. bovis* genome, complicating the curing process. However, using our curing process we typically attempted to cure 3-5 clones, which yielded 1-2 cured mutants. Such integration events have previously been reported in several mollicutes and exploited for mutant generation by homologous recombination (HR) ^40–42^. To limit this phenomenon, we employed a plasmid carrying a minimal heterologous *oriC* from *M. agalactiae* instead of that of *M. bovis* ^29^. Sequence divergence and size reduction were expected to reduce HR-mediated plasmid integration at the replication origin. Although rare, these events were still detected, making it preferable to screen multiple mutants to ensure successful curing. The incorporation of a counter-selection system could also accelerate plasmid curing as this approach has been successfully applied in *Pseudomonas putida* using the *sacB* gene ^32^. However no robust counter-selection markers currently exist for mollicutes, apart from CRISPR-based systems^13^.

This wet-laboratory work is complemented by the development of DTI, a design algorithm that facilitates tool selection depending on the targeted CDS and genome. Numerous gRNA design tools have been created since the emergence of CRISPR-based mutagenesis approaches, including some specifically for base editors ^43,44^. However, most are tailored to model organisms with pre-integrated reference genomes. By contrast, DTI is designed to incorporate the full genome of any strain or clone alongside that of the target gene(s), providing a comprehensive overview of all possible targets for our CRISPR-BE and CRISPR-BE VQR tools. Application of DTI to complete genomes of multiple mycoplasmas of human and veterinary interest showed that the VQR variant could potentially target more genes than the original version, and that combining both tools could allow targeted inactivation of over 90% of genes in most mycoplasmas. Implementation of these CRISPR-BE tools, including the VQR variant, in other mollicutes is highly feasible, provided transformation and a delivery vector (transposon or plasmid) are available. As demonstrated here, a mix-and-match approach can be followed to modify existing vectors with the genetic components described herein. Although transformation remains challenging and inefficient in some species, the high efficiency of these new tools means that, ultimately, a single transformant may suffice to obtain the desired mutant, unlocking remarkable opportunities to accelerate the study of these minimal pathogenic bacteria.

Overall, these CRISPR-BE tools markedly expand the genome engineering capacity of mycoplasmas by enabling broader targetability, efficient multiplex editing, and rapid, scarless mutant generation. Coupled with the DTI design algorithm, this platform provides a powerful and versatile framework for functional genomics in mollicutes. These advances are expected to accelerate the dissection of host–pathogen interactions and support the development of innovative control strategies.

## MATERIAL AND METHODS

### Oligonucleotides and plasmids

All oligonucleotides used in this study were supplied by Eurogentec and are described in Table S1. The plasmids constructed and used are described and listed in Table S2.

### Molecular cloning and plasmid generation

All following described plasmids were built via Gibson assembly using the NEBuilder HiFi DNA Assembly Cloning Kit (NEB, E5520S) used to assemble the DNA fragments and create the circular plasmids. Firstly, desired fragments were amplified with the Q5 DNA polymerase (NEB, M0491S). Once amplified, the PCR products were digested using the DpnI restriction enzyme (NEB, R0176S) following the manufacturer’s recommendations. The amplicons were then purified using the GFX PCR DNA and Gel Band Purification Kit (Cytiva) and eluted in 50 µL H2O. Following assembly, chemically competent *E. coli* NEB 5α cells (NEB, C2987H) were transformed with 2 μL of the assembly reaction and plated on 50 µg.mL^-1^ kanamycin LB agar plates, and incubated overnight at 37°C. Screening of the transformants was performed by first isolating their plasmid DNA using the NucleoSpin Plasmid kit (Macherey-Nagel), followed by enzymatic digestion of the DNA and restriction fragments size analysis by agarose gel electrophoresis. A subsequent screening step was performed by Sanger sequencing of the desired locus (Genewiz). Finally, whole plasmid sequencing was conducted through the use of the Plasmid-EZ service (Genewiz) or the Plasmidsaurus sequencing platform.

### Target sequence selection and sgRNA cassette construction

After a gene of interest was selected for inactivation, potential targetable loci were defined based on the possibility to generate a cytosine deamination in the editable window upstream of a PAM site. Once a target locus was selected, the corresponding target sequence (the 20 bp upstream of the PAM) was introduced into the gRNA expression cassette as follows. The 20-nucleotide target sequence was designed as complementary oligonucleotides F and R. Both primers were mixed in a hybridization buffer (20 mM Tris-HCl pH 7.5; 200 mM NaCl; 2 mM EDTA; 2 mM DTT), using 10 µL of each oligonucleotide at 100 µM and 20 µL of the hybridization buffer. Hybridization was conducted through heating at 95°C for 10 minutes then lowering the temperature at 0.1°C/second down to 20°C. In parallel, CRISPR-BE plasmids were cleaved using BseRI (NEB, R0581S) for 1h at 37°C following manufacturer’s instructions. The linearized product was subsequently purified using GFX PCR DNA and Gel Band Purification Kit.

After purification, target sequences and linearized plasmids were ligated using T4 DNA ligase (Thermofisher) at 20°C for 1h. NEB 5α competent *E. coli* cells (NEB, C2987H,) were then transformed with 2 μL of the ligation mix. Screening of the transformants was performed by first isolating their plasmid DNA using the NucleoSpin Plasmid kit (Macherey-Nagel), followed by enzymatic digestion with BseRI, as a negative control, and AgeI (NEB, R3552S), and restriction fragments size analysis by agarose gel electrophoresis. A subsequent screening step was performed by Sanger sequencing of the targeted locus using various oligonucleotide primers (Seq-End-TetR-F-FR/ Seq/begin-SpyoCas9-R-FR/Seq-PrePGenta-R-FR (Genewiz).

#### pFRIT4.0-DgRNA plasmid

The double gRNA design used in the generation of the pMT85_SpdCas9-dgRNA-pmcDA1-GentaR (pFRIT4.0-DgRNA) plasmids was based on pre-existing single gRNA plasmids. The first fragment was generated using a CRISPR-BE plasmid containing a gRNA target sequence, and was amplified using the primer pair secondgRNA-[pFRIT]-F and secondgRNA-[pFRIT]-R. The second gRNA expression cassette including the spiralin promoter^45^, the 20 bp target sequence or the “empty” target sequence site including the BseRI restriction sites, and a fibril terminator was PCR-amplified using the primer pair pFRIT-[secondgRNA]-F and pFRIT-[secondgRNA]-R. The two generated fragments were then assembled through Gibson assembly and following the manufacturer recommendations.

#### SpdCas9-VQR variants plasmids

The generation of the SpdCas9-VQR variants recognising the NGA PAM sequence required the substitution of three specific nucleotide bases in the coding sequence of SpdCas9: A3403T (causing D1135V), A4001C (causing R1335Q), C4009G (causing T1337R). The construction of these plasmids was carried out by Gibson Assembly using pFRIT4.0 (to create pFRIT4.0-VQR) as DNA templates. Two primers pairs (A1-pFRIT-SpyCas9QC-VQR-F-PH/A2-pFRIT-SpyCas9QC-VQR-R-PH and B1-pFRIT-SpyoCas9QC-VQR-F-PH/B2-pFRIT-SpyoCas9QC-VQR-R-PH) carrying the desired mutations were used to amplify two fragments from each plasmid by PCR. The assembly method then followed the described protocol above.

#### pFrHog replicative CRISPR-BE plasmids

These constructs were generated using the previously constructed pFRIT4.0 and pFRIT-VQR plasmids as DNA templates. The CRISPR-BE sequences, including the gRNA insertion site, SpdCas9-pmcDA1, and the Kana/GentaR resistance gene, were amplified using the primer pair pFrHog_[SpyoCas_genta]_F/pFrHog_[SpyoCas_genta]_R. A second fragment, containing the *M. agalactiae* origin of replication and the ColE1 origin, was amplified from the p20-1miniO/T plasmid^30,31^ using the primer pair pFrHog[pMaga_colE1]_F/pFrHog[pMaga_colE1]_R. Assembly and quality verification were performed as previously described.

#### pRES plasmid

The pRES plasmid was constructed using the PCJ15 plasmid^21^ as template. PCR amplification of the tetracycline resistance gene and the TnpR resolvase encoding gene along with its spiralin promoter was conducted using the primer pair pOriMaga-[pCJ15]-F -PH/ pOriMaga-[pCJ15]-R -PH. The chromosomal replicative origin of *M. agalactiae* was amplified with the p20-1miniO/T plasmid^30,31^ as a template using primers pCJ15-[pOriMaga]-F-PH/ pCJ15-[pOriMaga]-R -PH. Gibson assembly was carried out following manufacturer’s instructions and as previously described.

### Bacterial strains and culture

*M. bovis* type strain PG45 (Tax ID: 289397) was cultivated in SP4b medium^46^. Phenol red was used as a pH indicator in the medium to monitor growth. Tetracycline at 5 µg mL^−1^ and gentamicin at 100 μg·mL^−1^ were used for the selection of mutants. The *Escherichia coli* NEB-5α strain (New England Biolabs) was used for plasmid propagation and was cultivated in Luria broth (ThermoFisher), with the addition of kanamycin at 50 μg·mL^−1^ for selection.

### Transformation of *M. bovis*

*M. bovis* was transformed using a previously described polyethylene glycol mediated protocol^47,48^. Briefly, *M. bovis* cells from late log-phase cultures (2 mL per transformation) were transformed using 2.5 to 10 µg of plasmid DNA. After transformation, *M. bovis* cells were resuspended in 1 mL of SP4b medium and incubated for 3 h a 37°C, then serially diluted in order to plate on SP4b-agar plates.

### Induction of CRISPR-BE systems and mutant analysis

After transformation of *M. bovis* with the CRISPR-BE plasmids, induction of the deaminase was performed in liquid cultures using anhydrotetracycline (Abcam, ab145350) (in EtOH 50%) at a final concentration of 0.5 µg·mL^−1^, added in SP4b (1 mL).

Post-induction screening was performed on pools of transformants or on isolated clones by PCR (using Q5 High-Fidelity DNA Polymerase (NEB) or Advantage HF 2 PCR kits (Takara) and Sanger sequencing of the amplicons. The presence of the edited bases was manually checked from the Sanger chromatograms and qualitatively assessed through the BEAT program^23^. PCR were performed using DNA obtained through a rapid DNA extraction method. Briefly, 200 µL of late-log grown liquid culture are centrifuged at 10,000×*g* for 10 min at room temperature. The pellet of cells is then resuspended in 100 µL of TE solution (Tris-HCL 10mM; p7.5; EDTA 1mM pH 7.5) before incubation at 95°C for 10 minutes for cell lysis. The mixture is used as a source of DNA for PCR.

### Whole genome sequencing

Genomic DNA of *M. bovis* was purified from 10 mL cultures, using the Nucleobond AXG 20 columns and kit (Macherey-Nagel) following the manufacturer’s instructions. Once purified and concentrated at 50 ng.µL^-1^, samples were sent to the Plasmidsaurus sequencing platform. Whole genome sequencing and assembly were performed by Plasmidsaurus using Oxford Nanopore Technology and Illumina sequencing. To do so, an amplification-free long-read library was constructed with ONT v14 library prep chemistry. Sequencing was performed using a primer-free protocol on R10.4.1 flow cells, with raw data delivered in fastq format. A minimum raw read Q-score of 10 (90% accuracy) was required. For hybrid ONT + Illumina sequencing, the long-read assembly was further polished with paired-end 2×150 bp Illumina reads.

The bioinformatics pipeline for assembly and polishing consisted of the following steps: first, base-calling was performed with Dorado v4.3 using default Q10 quality filtering, followed by removal of the lowest-quality 5% of reads with Filtlong v0.2.1. Reads were initially down-sampled to 250 Mb to generate a rough assembly with Miniasm v0.3, then re-down-sampled to ∼100× coverage while giving heavy weight to high-quality reads to retain small plasmids. Genome assembly was then conducted with Flye v2.9.1, and the resulting assembly was polished using Medaka v1.8.0. Assemblies were annotated with Bakta v1.6.1, contigs analysed with Bandage v0.8.1, and genome completeness and contamination assessed with CheckM v1.2.2. For hybrid sequencing, the ONT assembly was additionally polished with Illumina reads using Polypolish v0.6.0, producing the final high-quality polished hybrid genome assembly.

### Resolution of integrated pMT85-based CRISPR-BE

The resolution of integrated pMT85/2res plasmid cassettes was performed through a secondary transformation with the constructed pRES plasmid (Fig. S5). The process was based on works conducted by Janis and colleagues^21^. The transformation of *M. bovis* was carried out as described above, followed by plating, colony isolation and successive passages in SP4b medium supplemented with tetracycline (5 µg·L⁻¹). PCR was used to confirm elimination of the cassette, using DNA extracted from late-log-phase mutant cultures. The PCR amplification was carried out using primers Seq-End-TetR-F-FR/ Seq-begin-SpyoCas9-R-FR targeting regions within the inserted CRISPR-BE cassette. Following confirmation of resolution, non-selective passages were performed, and a final PCR screening targeting the pRES plasmid was conducted to verify curing of the pRES plasmid. The primer pairs Seq-beginTetM-R/MT85-after-TetM and pGenta-F/pGenta-R were used to amplify a region of the pRES plasmid. Mutants were also cultured in parallel in gentamicin (100 µg·mL⁻¹) and tetracycline (5 µg·mL⁻¹) when resolution and curing were anticipated to have occurred as a second screening process.

### Deaminase Target Identifier (DTI)

The Deaminase Target Identifier package is a Python-based tool that was developed to identify possible targets of CRISPR-BE tools in a set of nucleotide sequences (Fig. 5). The tool identifies target codons of interest for the generation of stop codons (CAA/CAG/TGA in the correct ORF) upstream of a PAM. The search for these sequences is customisable based on user-defined parameters. Briefly, the DTI requires a file containing the set of CDS of interest in a fasta format (.fasta). The customisable parameters are as follows: the choice of research codon (CAA/CAG/TGA or other non-stop generating codons); pams: Protospacer Adjacent Motifs (PAM); deamination window: window of search upstream from the PAM. The output is a text result file created in the folder containing the CDS and genome files. For each CDS, the sequences of nucleotides identified as possible targets are listed. All relevant information is presented in the following GitHub page: https://github.com/MarcDcls/DTI.

## Abbreviations

CDS: coding sequence
CRISPR: clustered regularly interspaced short palindromic repeat
CRISPR-BE: CRISPR Base Editor
IR: inverted repeat
SpCas9: *Streptococcus pyogenes* Cas9
IS: insertion sequence
PAM: protospacer adjacent motif
PI: PAM interacting domain.

## Acknowledgements

We are thankful to Marie Jittasevi and Julien Vadé for technical help in bacterial cultures and plasmid constructions. We also thank the RAMbo-V consortium for much appreciated discussions on the work and manuscript.

## Author contributions

P.H.: conceptualization, methodology, validation, formal analysis, data curation, investigation, writing – original draft, writing – review and editing; M.C.: software, formal analysis; T.I.: methodology ; C.L.: conceptualization, methodology, formal analysis, supervision; G.G.: investigation, formal analysis; A.B: conceptualization, supervision; E.B.: conceptualization, writing – review and editing; L.B.: conceptualization, methodology, formal analysis, supervision; Y.A.: conceptualization, methodology, formal analysis, investigation, data curation, supervision, writing – original draft, writing – review and editing; P.S.-P.: conceptualization, methodology, formal analysis, data curation, supervision, writing – original draft, writing – review and editing; F.R.: conceptualization, methodology, formal analysis, data curation, supervision, writing – original draft, writing – review and editing.

## SUPPLEMENTARY FIGURES

**Figure S1:**
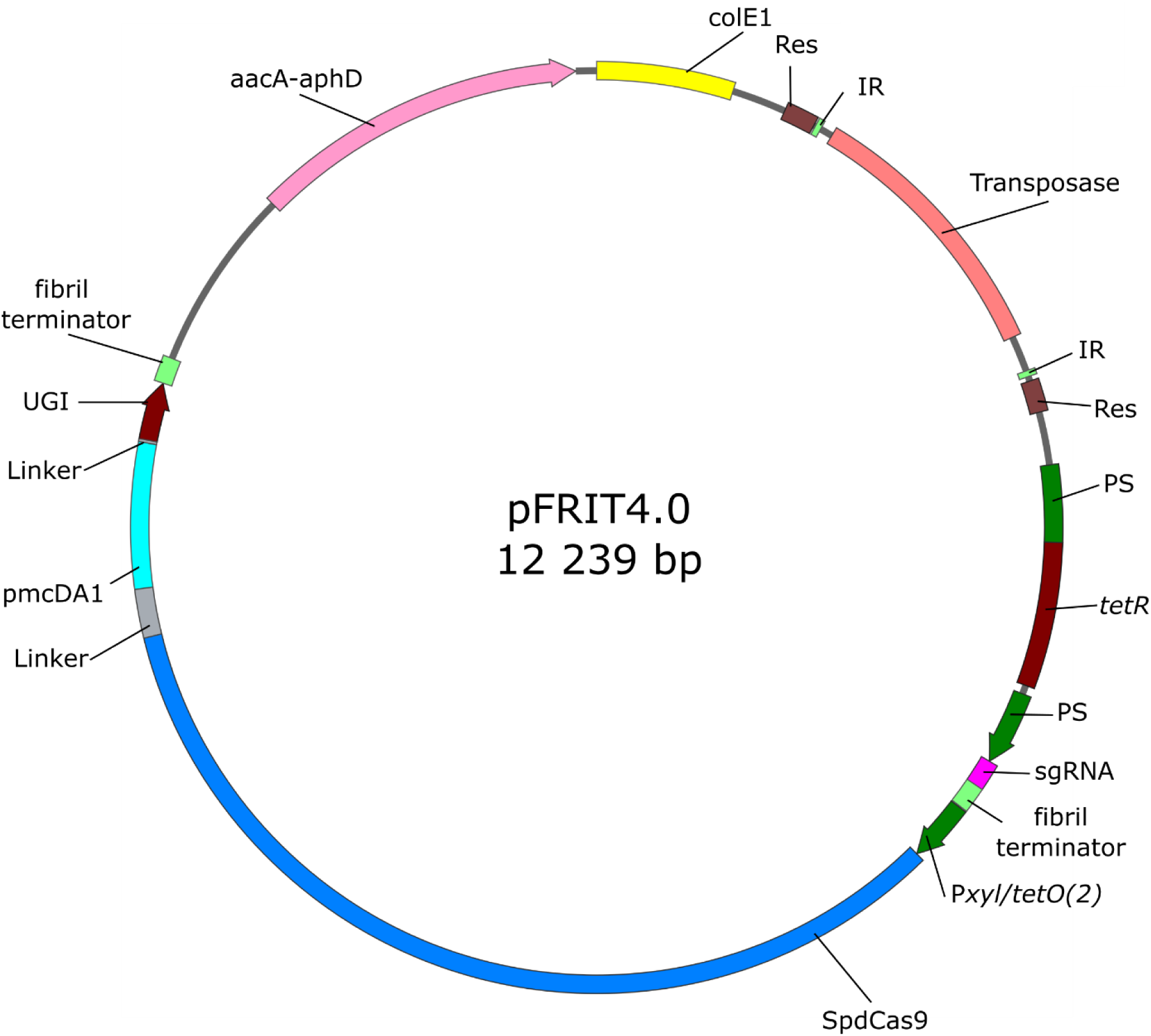
Original CRISPR-BE tool plasmid used in Mycoplasmas. ColE1, replicative origin from *E. coli;* TnpA, transposase; Res, resolvase sequences allowing for resolution; IR, inverted repeats; PS, promoter of the *S. citri* spiralin gene; *tetR*, repressor gene ; sgRNA controlled by PS promoter and terminator from the *S. citri* fibril gene; P*xyl/tetO(2*) inducible promoter that controls the expression of the fusion protein SpdCas9-pmcDA1-UGI; SpdCas9 with position of the codons corresponding to the modified amino acids indicated; aacA-aphD, gentamicin resistance marker.

**Figure S2.**
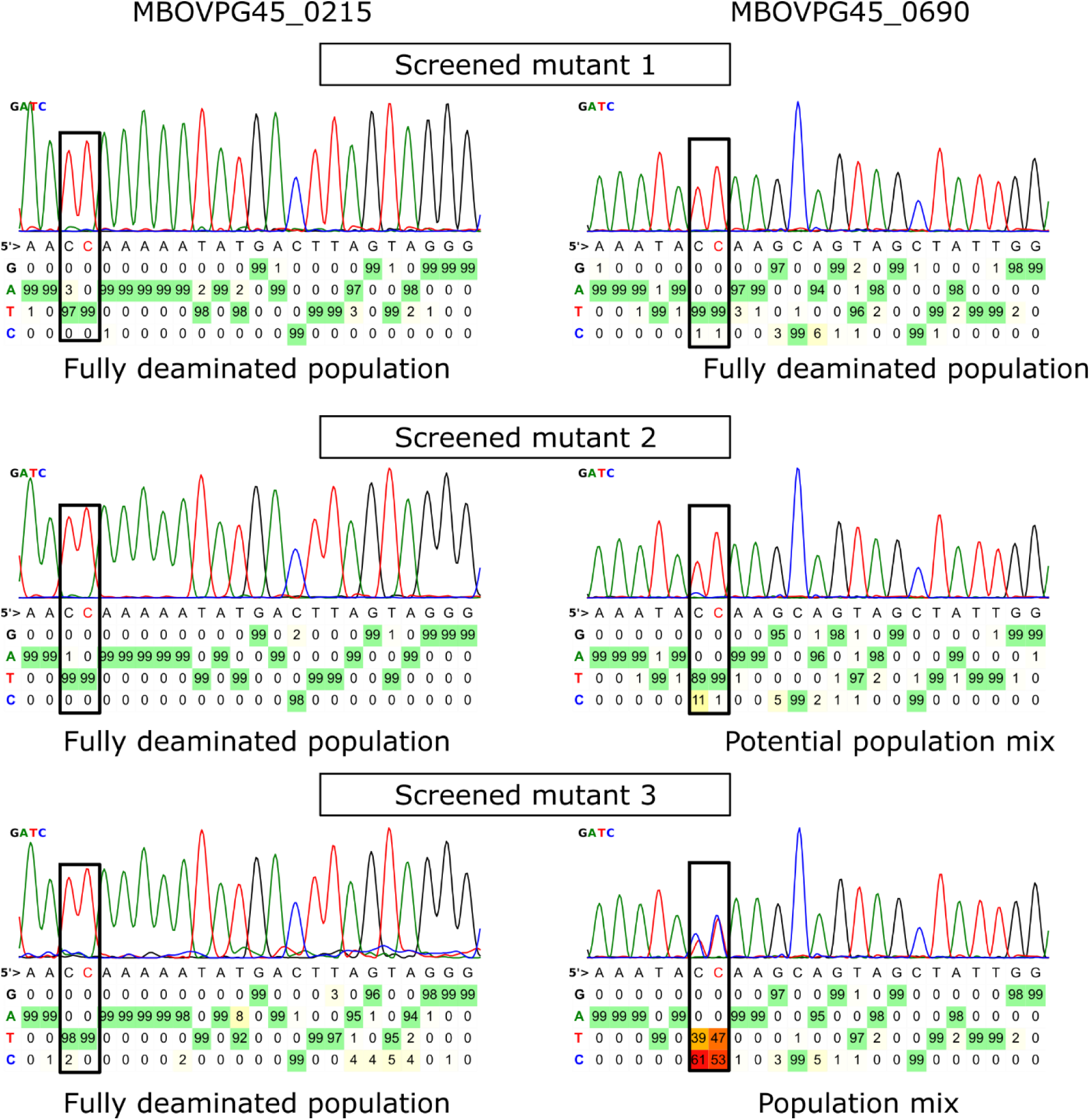
Mutant screening of dual target deamination. Screening of mutants obtained for targeting MBOVPG45_0215 and MBOVPG45_0690 was first conducted by Sanger sequencing of eight isolated clones. Following analysis, three displayed deaminated cytosine residues at the targeted region of MBOVPG45_0215 (Left). These three validated mutants were then screened by Sanger sequencing of the second target region of MBOVPG45_0690. Analysis revealed varying degrees of cytosine deamination at the second target site. Mutant 1 exhibited an apparently fully deaminated profile. Mutant 2 showed a minor background signal of cytosine base calling at one of the two targeted cytosines. Finally, in contrast, mutant 3 appeared as a mixed population of cells, with the targeted cytosines still showing strong base calling for the original cytosine (C).

**Figure S3.**
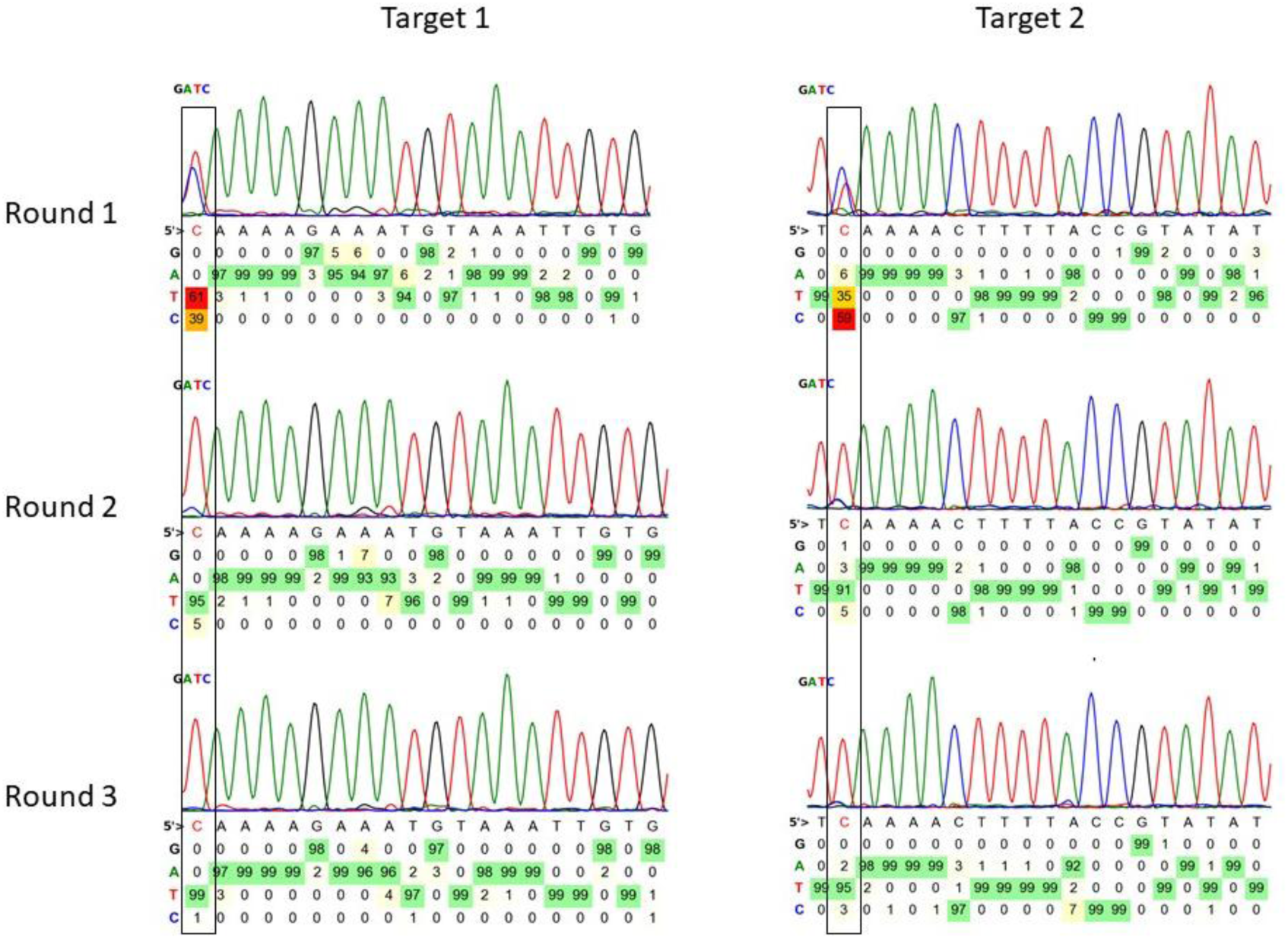
Multiple induction rounds for knockout of *ISMbov1* genes. To obtain full deamination on all *ISMbov1* coding sequences, three rounds of induction were necessary. An example of the profiles obtained after one, two or three rounds of deamination for one of the three clones is presented here. The first round was conducted as previous cultures were, with overnight induction, followed by gDNA extraction and Sanger screening. After primary analysis, which revealed that full deamination had not occurred, two supplementary passages under inducting conditions were performed and gDNA was extracted at each of them for PCR amplification and Sanger sequencing.

**Figure S4.**
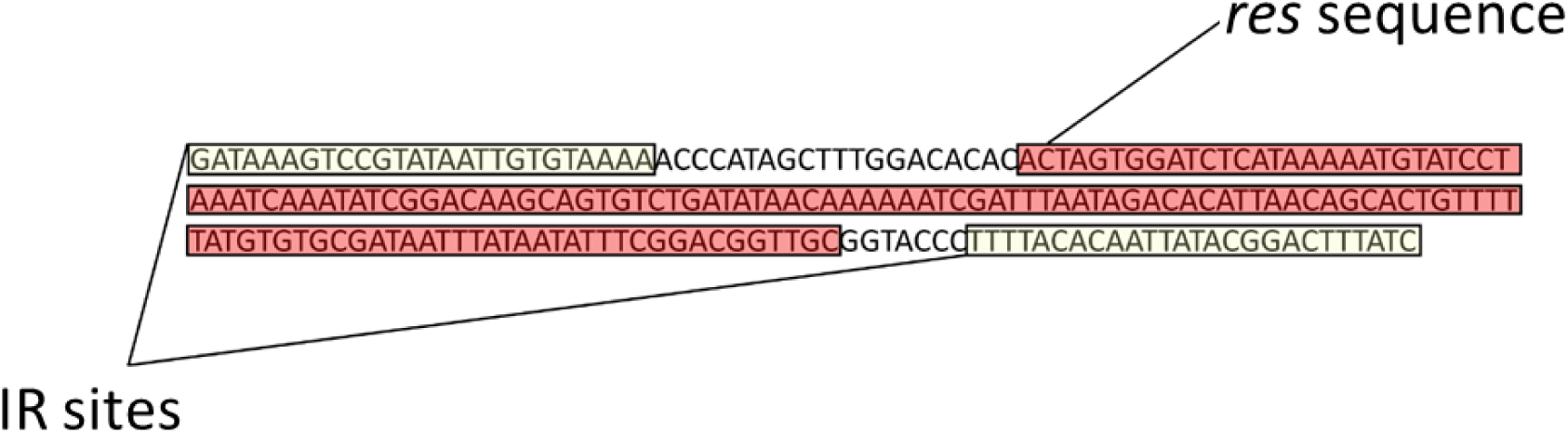
Genome structure after resolution of the transposon-based CRISPR-BE. After recombination between the *res* sequences induced by the *γδ* resolvase, a 218 bp scar remains at the site of integration of the transposon. It includes the remaining *res* sequence (coloured in red) and inverted repeats (IR) at the extremities of the transposon (coloured in green).

**Figure S5.**
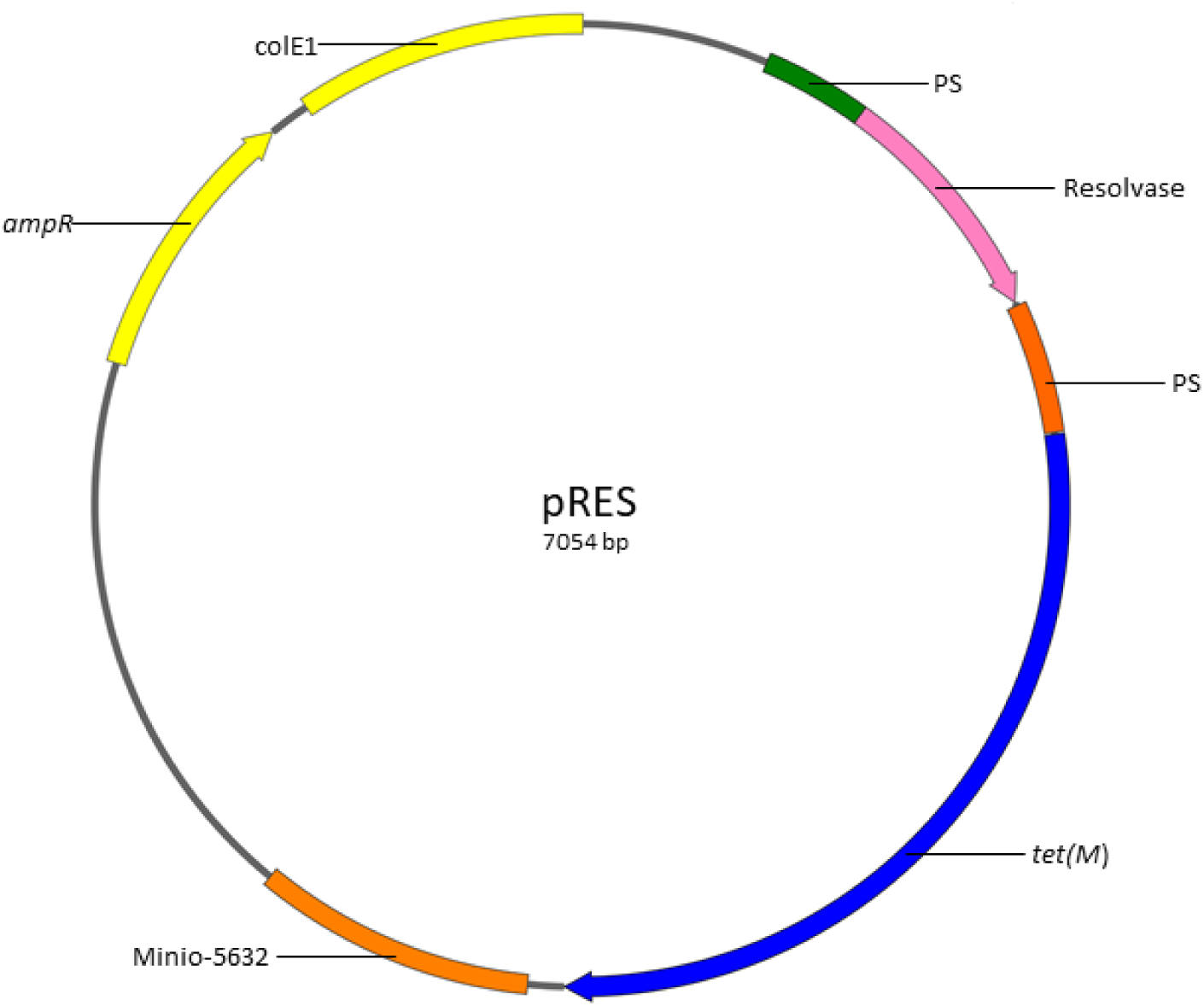
Map of pRES plasmid. PS, promoter of the spiralin gene; resolvase, gene encoding the *γδ* resolvase; *tet(M)* tetracycline resistance marker; Minio-5632, Minimal *oriC* region from *M. agalactiae* 5632 genome; *ampR*, ampicillin resistance marker; ColE1, replicative origin from *E. coli*.

**Figure S6.**
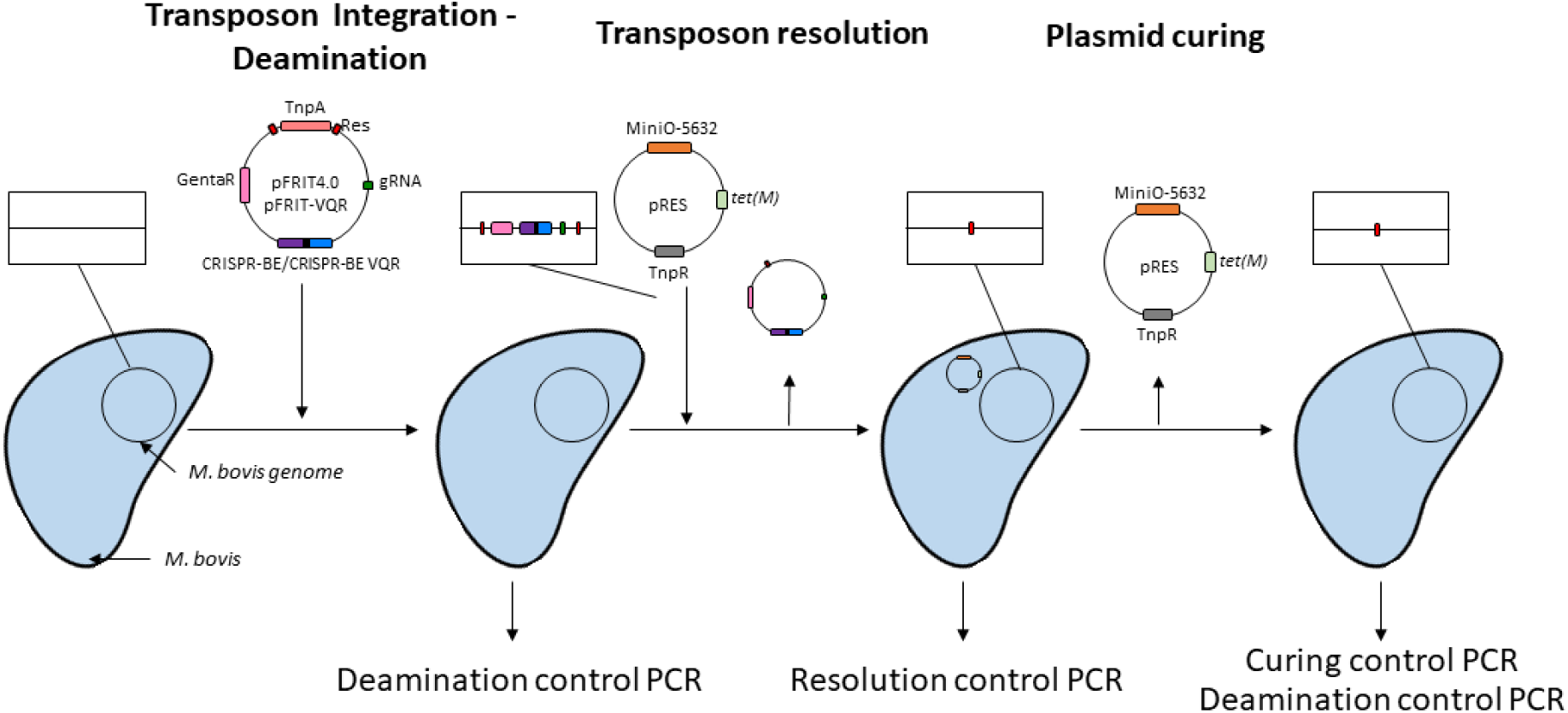
Workflow for generation of deaminated-resolved-cured mutants with a transposon-based tool. The multi-step process for the generation of deaminated-resolved-cured (DRC) mutants involves first the transformation of the mycoplasma, the integration of the transposon harbouring the CRISPR-BE and induction of its expression to obtain a deaminated mutant. PCR and Sanger sequencing screening is then used to select a deaminated mutant. Resolution of the transposon is then achieved by transformation with the pRES plasmid. Continued passages of isolated mutants under selection with tetracycline 5 µg.µL^-1^ are combined with DNA extraction and PCR amplification at each passage to verify the excision of the transposon cassette. Once excision is validated by PCR, gentamicin susceptibility is confirmed by culture in gentamicin selective media at 100 µg.mL^-1^. Following confirmation, the isolated mutants are cultivated in non-selective media and subcloned until negative PCR amplification of a pRES region and inability to grow in selective tetracycline medium are obtained. Final verifications were done through Sanger sequencing of the deamination site and WGS to verify genomic scar presence.

**Figure S7.**
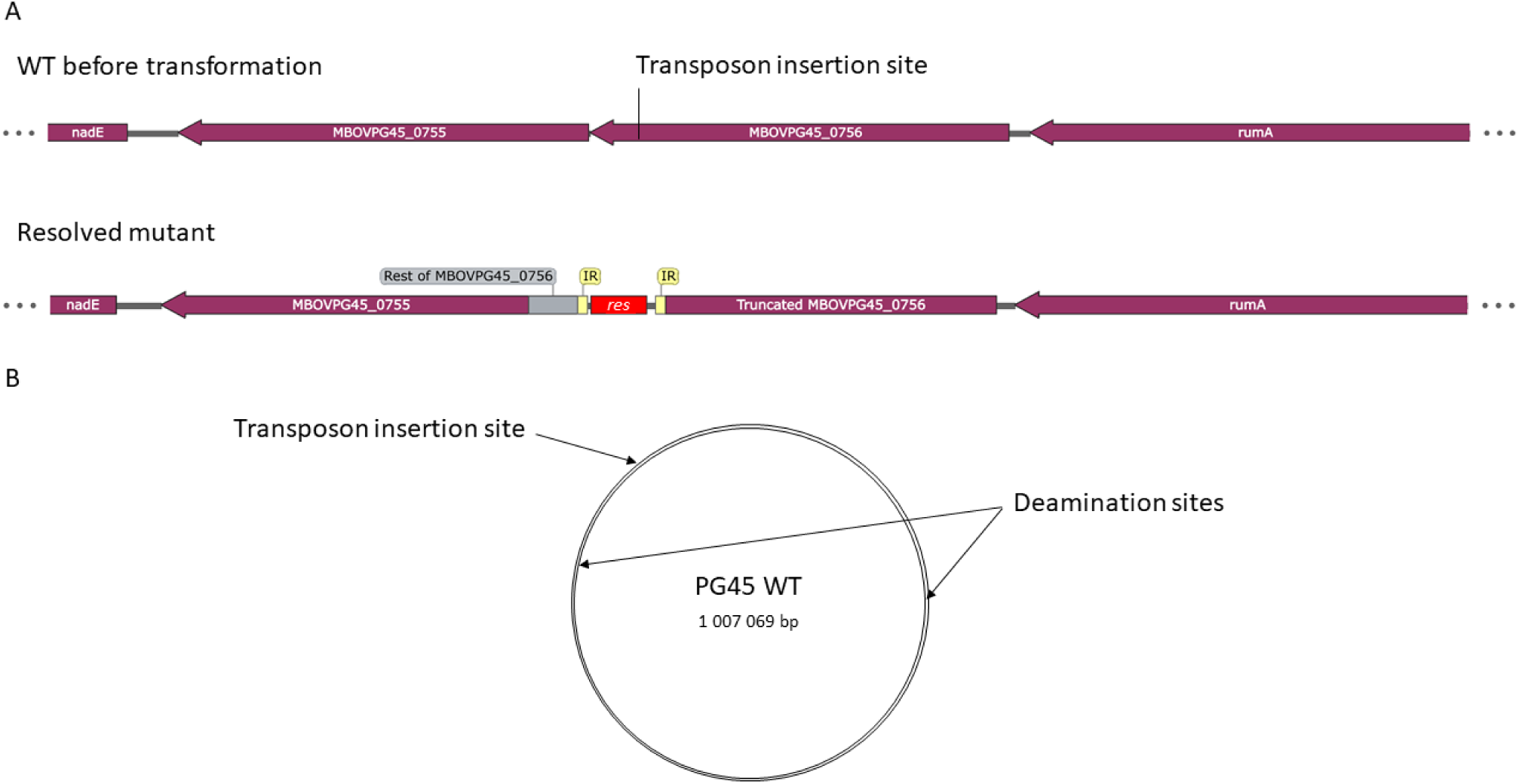
Genome structure at the insertion site of the transposon within the double MBOVPG45_0215 and MBOVPG45_0690 mutant. WGS sequencing revealed the successful resolution of the transposon and indicated insertion locus of the employed CRISPR-BE was intragenic to the MBOVPG45_756 coding sequence. (A) Genome structure in the WT and in the resolved mutant. (B) Localisation of the transposon insertion site and targeted sites in the genome of *M. bovis* PG45.

**Figure S8.**
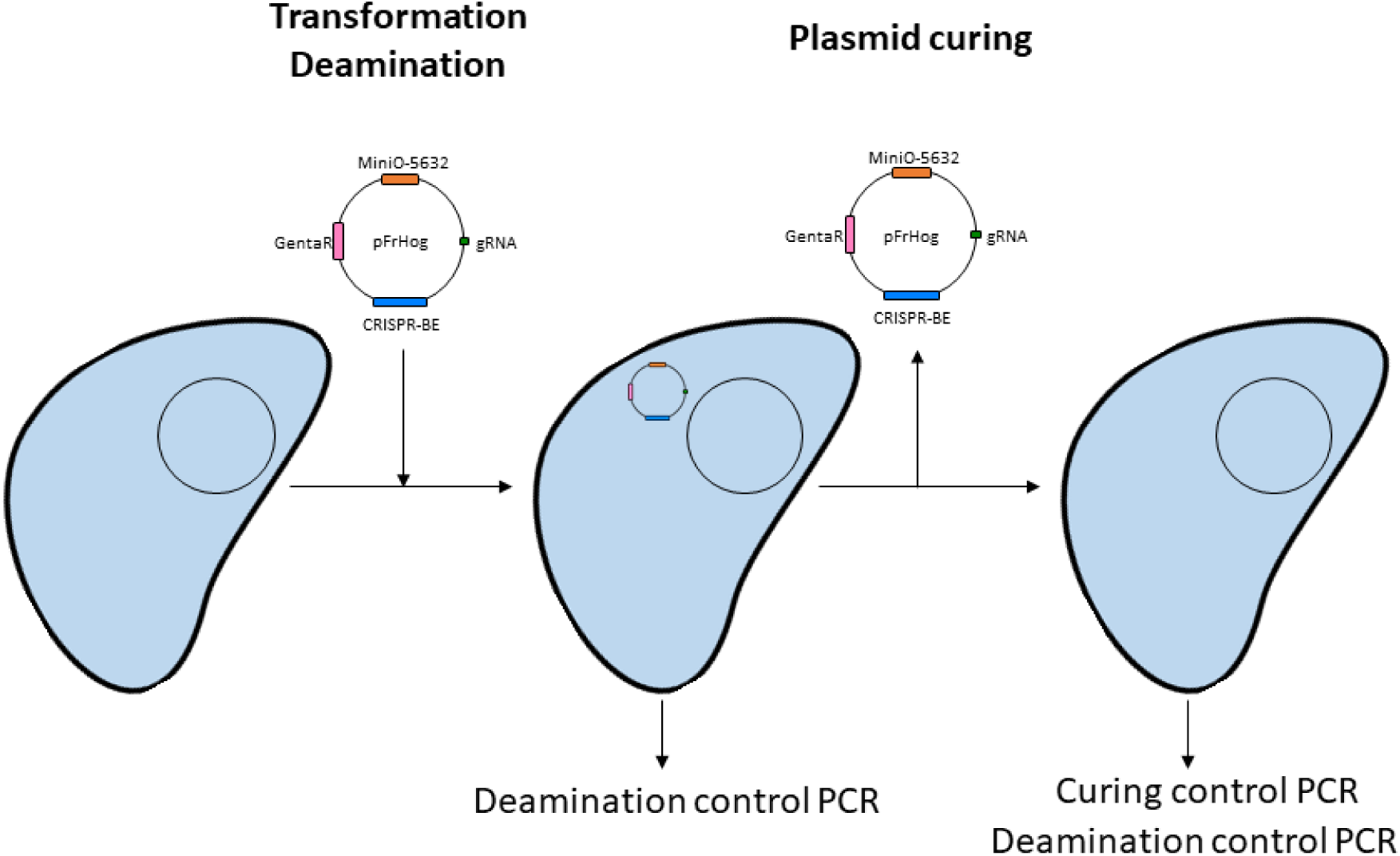
Workflow for generation of deaminated-resolved-cured mutants with a replicative plasmid-based tool. In the case of the obtained MBOVPG45_0215 mutant with the pFrHog plasmid, the process to obtain a cured mutant is a condensed version of the previously described DRC pipeline. After transformation, a single round of induction is necessary to obtain the desired base-editing. PCR verifications for the presence of mutation and cultures in selective medium are used to follow the process. Curing of the plasmid is then obtained by passages in non-selective medium. Once both validation steps are validated, the isolated mutant is considered to be cured and can be reverified at the deamination site through PCR and Sanger sequencing.

**Figure S9.**
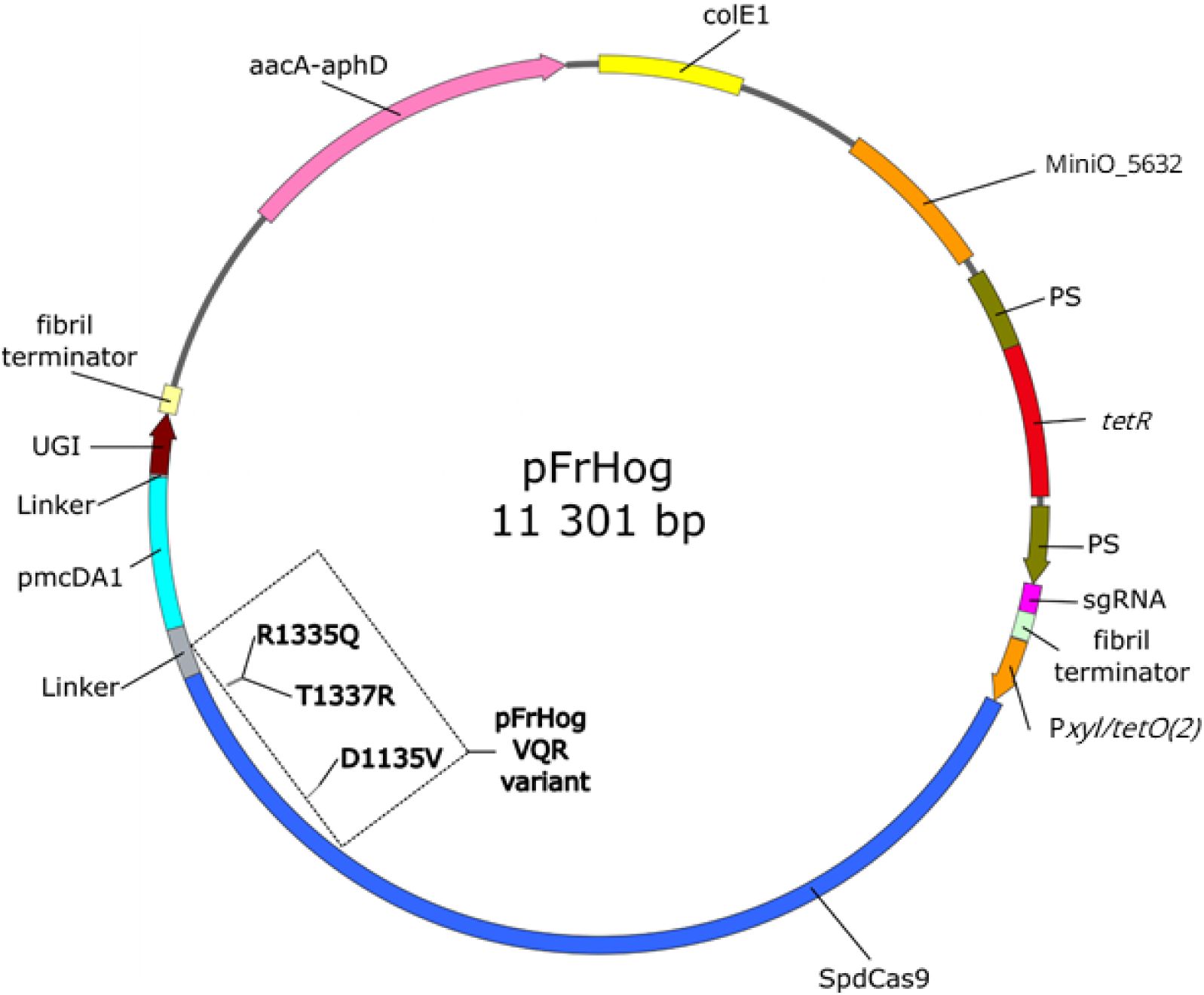
Map of pFrHog and pFrHog VQR variant plasmids. Both CRISPR-BE plasmids have the same structure except the VQR modifications in SpdCas9 of the VQR variant. ColE1, replicative origin from *E. coli;* Minio-5632, minimal *oriC* region from *M. agalactiae* 5632 genome; PS, promoter of the *S. citri* spiralin gene; *tetR*, repressor gene; sgRNA controlled by PS promoter and terminator from the *S. citri* fibril gene; P*xyl/tetO(2*) inducible promoter that controls the expression of the fusion protein SpdCas9-pmcDA1-UGI; SpdCas9 with position of the codons corresponding to the modified amino acids indicated; aacA-aphD, gentamicin resistance marker.

**Figure S10.**
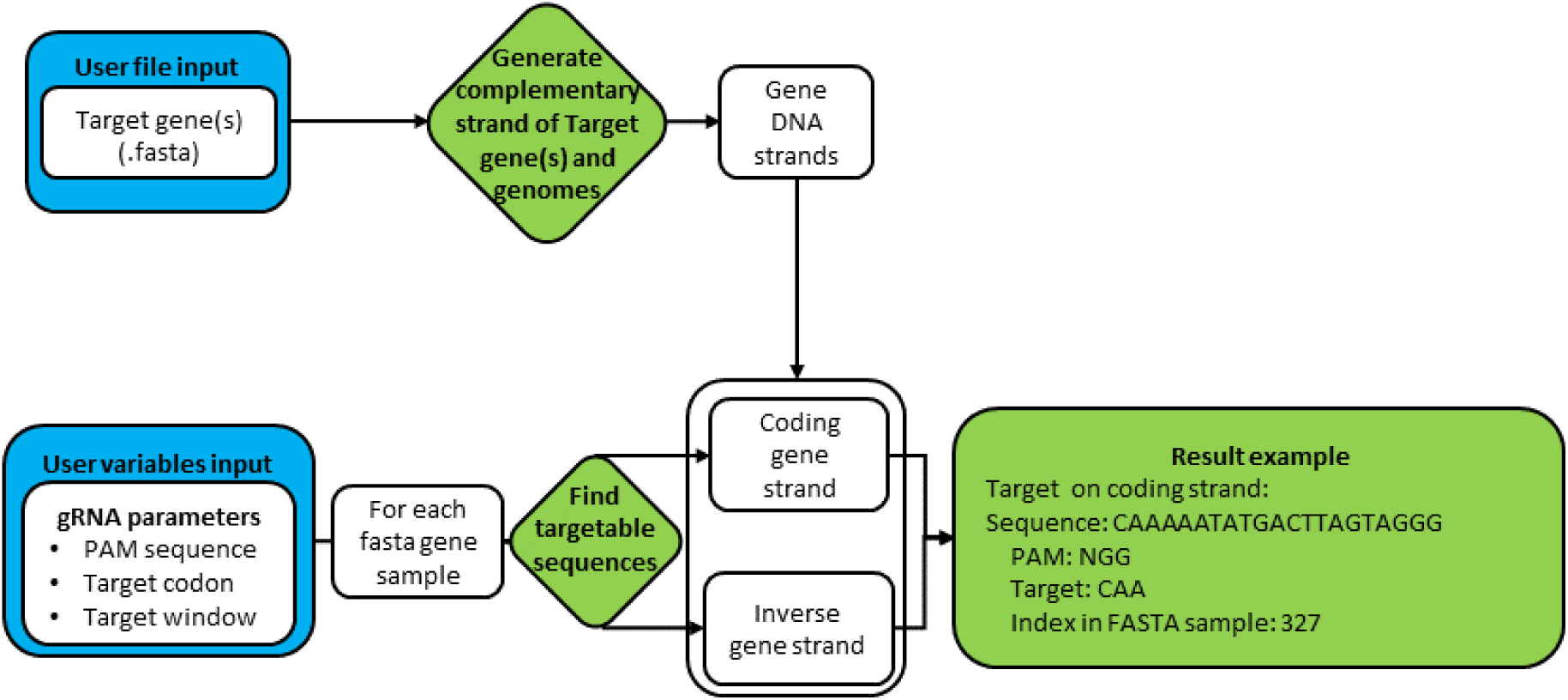
Diagram of the Python-based algorithm Deaminase Target Identifier (DTI). The pipeline is used to identify targetable regions within the predicted coding sequences of a bacterial genome.

